# Collection of Biospecimens from the Inspiration4 Mission Establishes the Standards for the Space Omics and Medical Atlas (SOMA)

**DOI:** 10.1101/2023.05.02.539108

**Authors:** Eliah G. Overbey, Krista Ryon, JangKeun Kim, Braden Tierney, Remi Klotz, Veronica Ortiz, Sean Mullane, Julian C. Schmidt, Matthew MacKay, Namita Damle, Deena Najjar, Irina Matei, Laura Patras, J. Sebastian Garcia Medina, Ashley Kleinman, Jeremy Wain Hirschberg, Jacqueline Proszynski, S. Anand Narayanan, Caleb M. Schmidt, Evan E. Afshin, Lucinda Innes, Mateo Mejia Saldarriaga, Michael A. Schmidt, Richard D. Granstein, Bader Shirah, Min Yu, David Lyden, Jaime Mateus, Christopher E. Mason

## Abstract

The SpaceX Inspiration4 mission provided a unique opportunity to study the impact of spaceflight on the human body. Biospecimen samples were collected from the crew at different stages of the mission, including before (L-92, L-44, L-3 days), during (FD1, FD2, FD3), and after (R+1, R+45, R+82, R+194 days) spaceflight, creating a longitudinal sample set. The collection process included samples such as venous blood, capillary dried blood spot cards, saliva, urine, stool, body swabs, capsule swabs, SpaceX Dragon capsule HEPA filter, and skin biopsies, which were processed to obtain aliquots of serum, plasma, extracellular vesicles, and peripheral blood mononuclear cells. All samples were then processed in clinical and research laboratories for optimal isolation and testing of DNA, RNA, proteins, metabolites, and other biomolecules. This paper describes the complete set of collected biospecimens, their processing steps, and long-term biobanking methods, which enable future molecular assays and testing. As such, this study details a robust framework for obtaining and preserving high-quality human, microbial, and environmental samples for aerospace medicine in the Space Omics and Medical Atlas (SOMA) initiative, which can also aid future experiments in human spaceflight and space biology.

## Introduction

Our human space exploration efforts are at a unique transition point in history, with more crewed launches and human presence in space than ever before^1^. We can attribute this to the commercial spaceflight sector entering an industrial renaissance, with multiple companies forming collaboration and competition networks to send commercial astronauts into space. This recent evolution of human space exploration endeavors presents a valuable opportunity to accumulate more biological research specimens and improve our understanding of the impact of spaceflight on human health. This is critical since there is still much to learn about the varied biological responses to the spaceflight environment, characterized by microgravity and space radiation landscape^2^. The impact of spaceflight on human health includes musculoskeletal deconditioning^3^, cardiovascular adaptations^4^, vision changes^5^, space motion sickness^6^, neurovestibular changes^7^, immune dysfunction^8^, and increased risk of rare cancers^9^, among other changes^2^. However, we are still at the very beginning of the work to catalog biological responses to spaceflight exposure at the molecular resolution.

Prior work has characterized molecular changes that occur during spaceflight in astronauts. These include changes in cytokine profiles^8, 10, 11^, urinary albumin abundance^12^, and hemolysis^13^. Furthermore, multi-omic assays have provided genomic maps of structural changes in DNA^14–16^, RNA expression profiles^11, 17, 18^, sample-wide protein measurements^17, 19, 20^, and metabolomic status^17^. Additionally, International Space Station (ISS) surfaces have been studied with longitudinal microbial profiles to track microbial pathogenicity and evolution to assess their potential influence on crew health^21, 22^. To better improve our understanding of both human and microbial biology in space, it is critical that these analyses continue and expand as more spacecraft and stations are built and flown.

Combining and comparing work from prior missions in these new spacecraft and stations is especially important to overcome the small sample sizes and highlights a need for standardization between missions. In addition, recruiting large cohorts of astronauts is difficult, as the ISS typically can only house up to six astronauts at a time. As of the time of writing, only 647 humans have been to space, starting with the launch of Yuri Gagarin in 1961. Studies have spanned the Vostok program, Project Mercury, the Voskhod program, Project Gemini, Project Apollo, the Soyuz program, the Salyut space stations, MIR, the Space Shuttle Program, SkyLab, Tiangong Space Station, and the ISS. From the breadth of experiments that have been performed on the ISS, only a minority have specifically been human research-oriented^23^, and just a subset involve omics studies. The NASA Twin Study created the most in-depth multi-omic study of astronauts prior to Inspiration4, but was limited to one astronaut and one ground control^17^. All of these factors have limited the statistical power of astronaut omic experiments and increase the difficulty of providing robust scientific conclusions. Standardizing biospecimen collections across multiple missions will create larger sample-sets needed to draw these conclusions.

Here, we establish the standard biospecimen sample collection and banking procedures for the Space Omics and Medical Atlas (SOMA). A key goal of SOMA is to standardize biospecimen collection and processing for spaceflight, to generate high-quality multi-omics data across spaceflight investigations. This paper provides sample collection methods built for standardized collections across different crews and missions. These can generate harmonized datasets with greater statistical power and thus increase our scientific return yields from spaceflight investigations. We also present metrics on sample collection yields, instances of prior astronaut sample collection in scientific literature, and considerations for improvement of sample collection on future missions based on crew feedback. In its inaugural use case, these samples were collected from the Inspiration4 (I4) astronaut cohort and are currently in use for several other missions (Polaris Dawn, Axiom-2), which will enable continued utilization for future crewed space missions.

## Results

### Biospecimen Collection Overview

We formulated and executed a sampling plan that spans a wide range of biospecimen samples: venous blood, capillary dried blood spots (DBSs), saliva, urine, stool, skin swabs, skin biopsies, and environmental swabs (**Fig 1a**). The collection of various types of samples covered the scope of previous assays on astronaut samples (**Table 1**), but also enabled newer omics technologies, such as spatially resolved, single-molecule, and single-cell assays.

**Figure 1:**
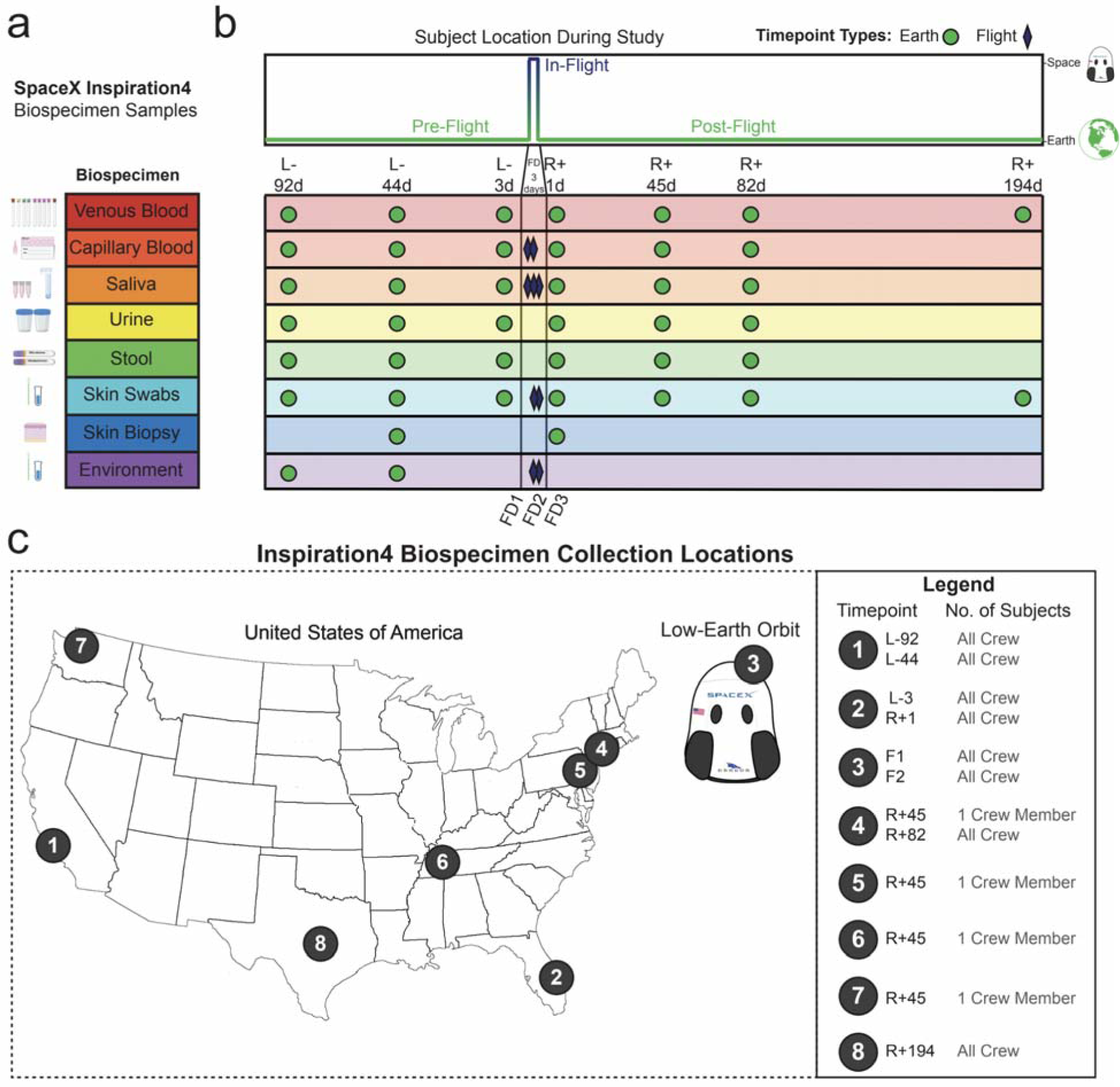
Biospecimen Samples and Collection Locations. **(a)** List of biospecimen samples collected over the course of the study. **(b)** Timepoints for each biospecimen sample collection. “L-” denotes the number of days prior to launch. “R+” denotes the number of days after return to Earth. “FD” denotes which day of the flight a sample was collected. **(c)** Location of each collection timepoint.

**Table 1:**
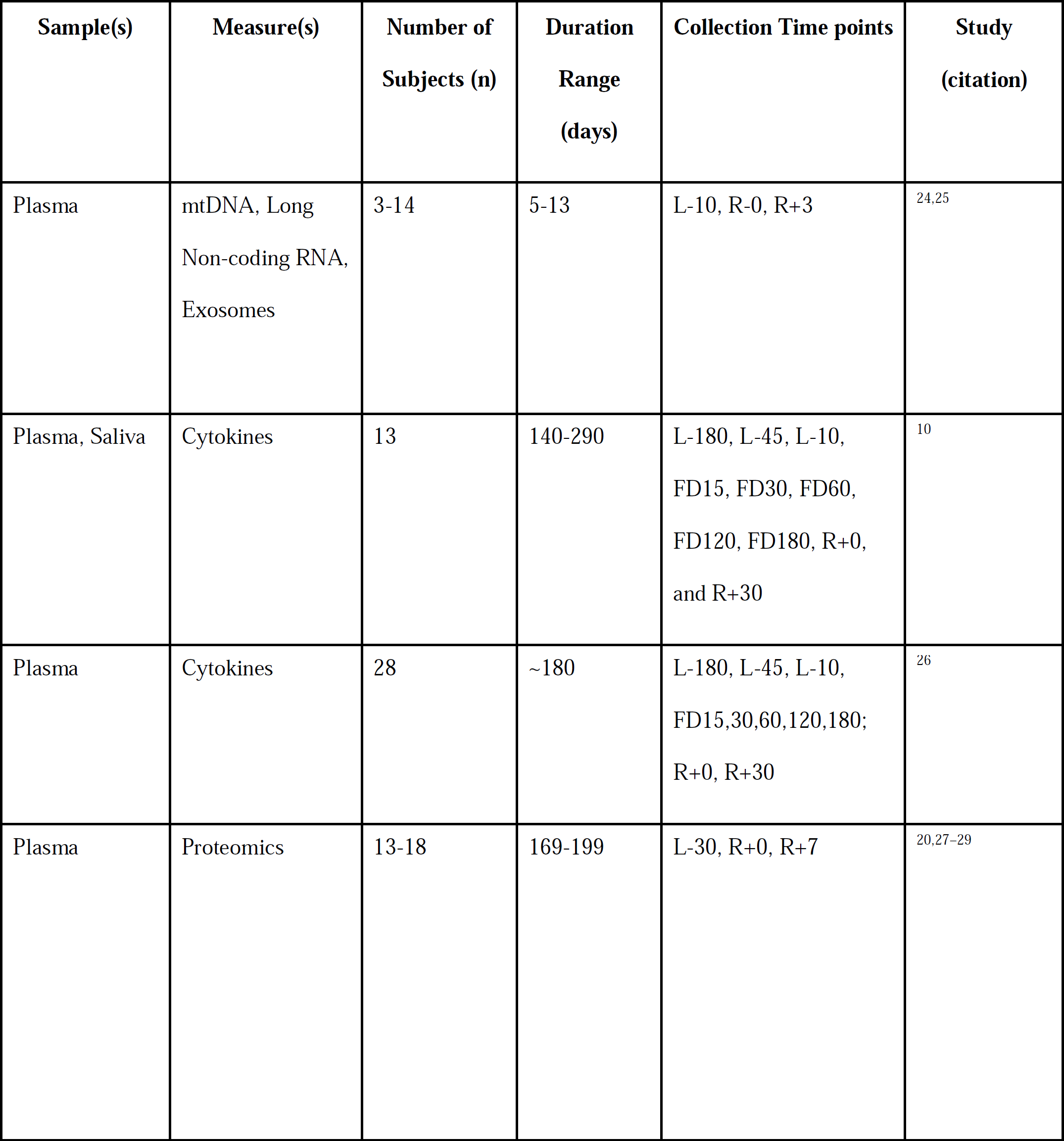
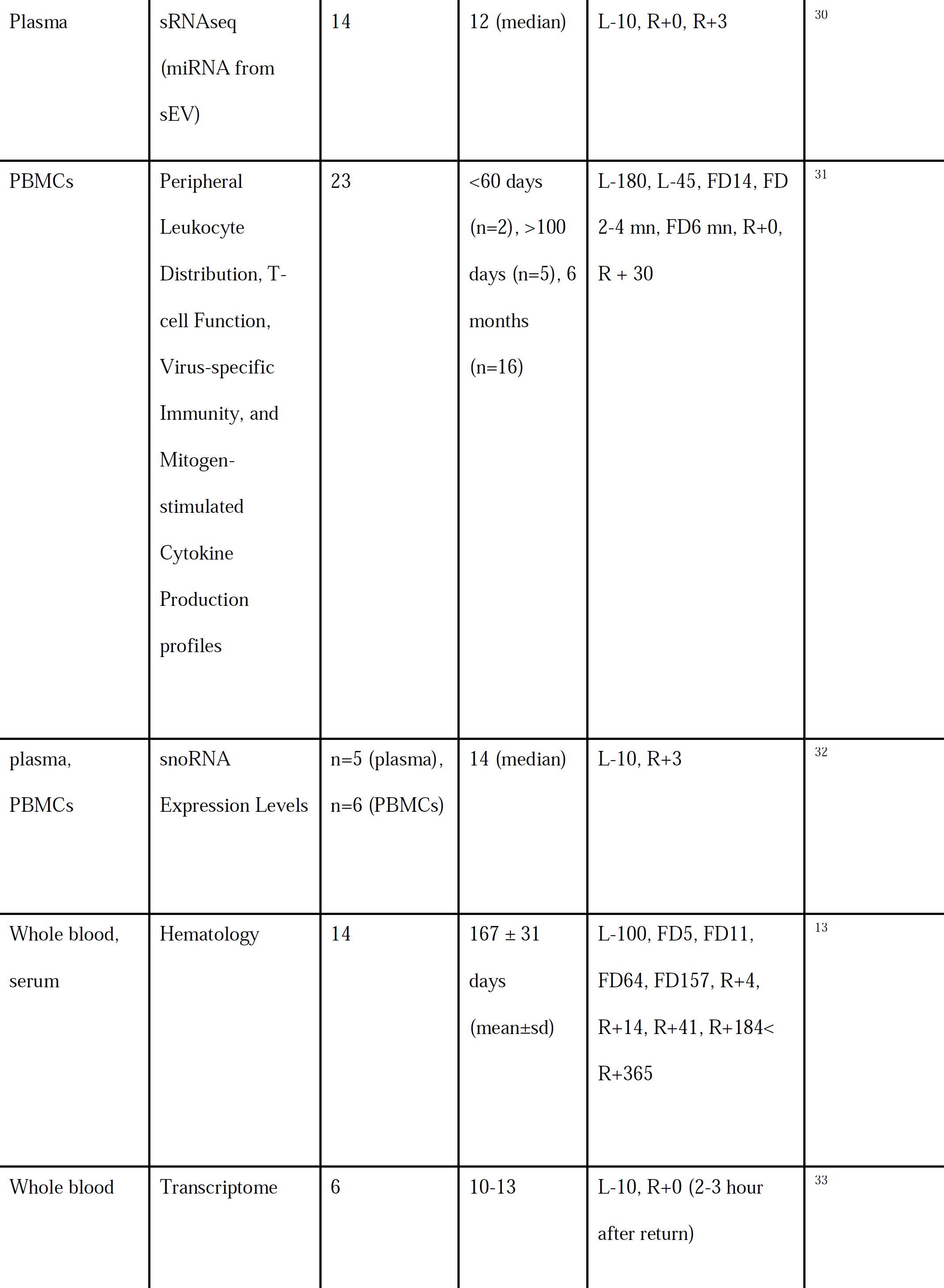
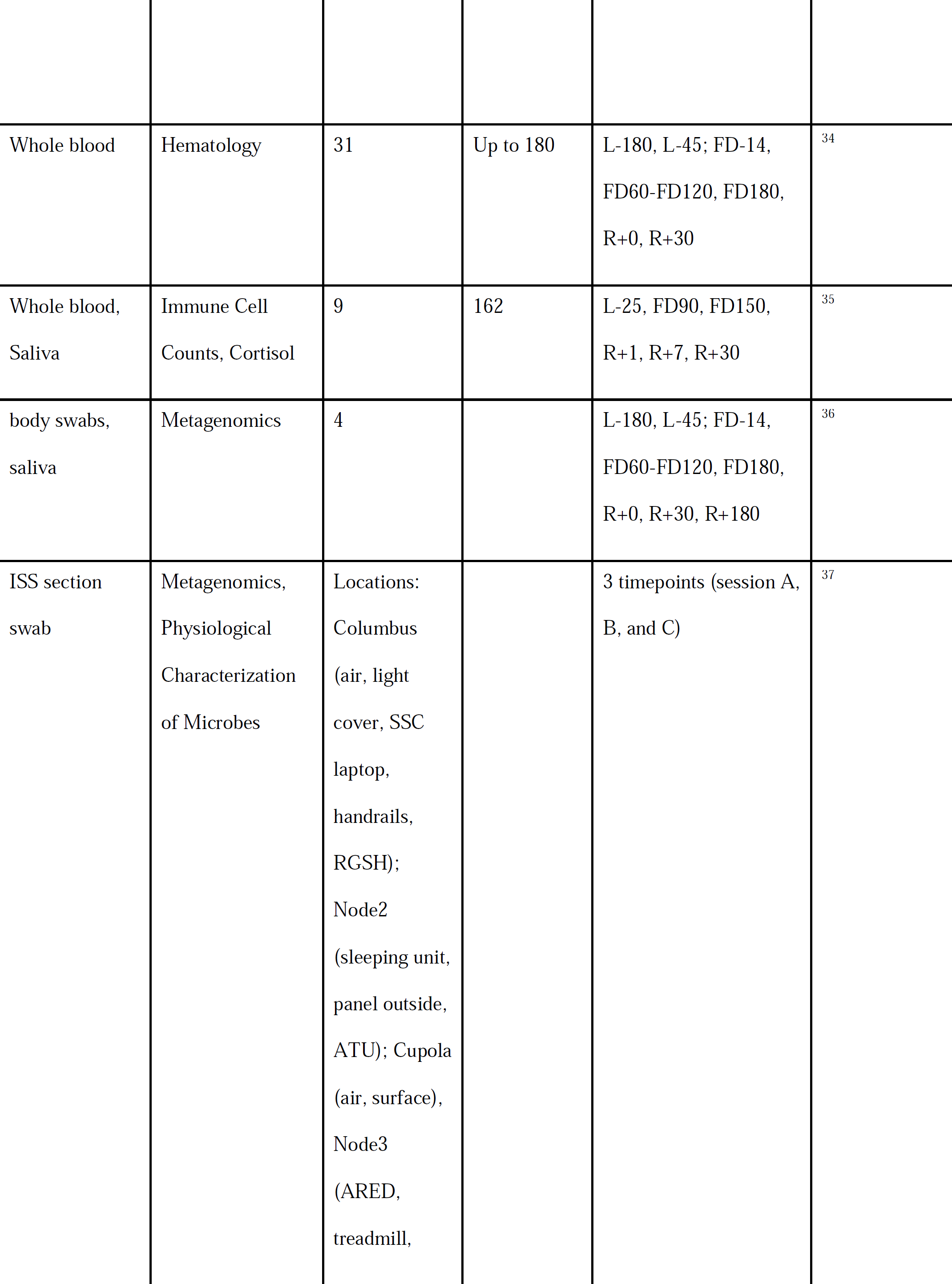
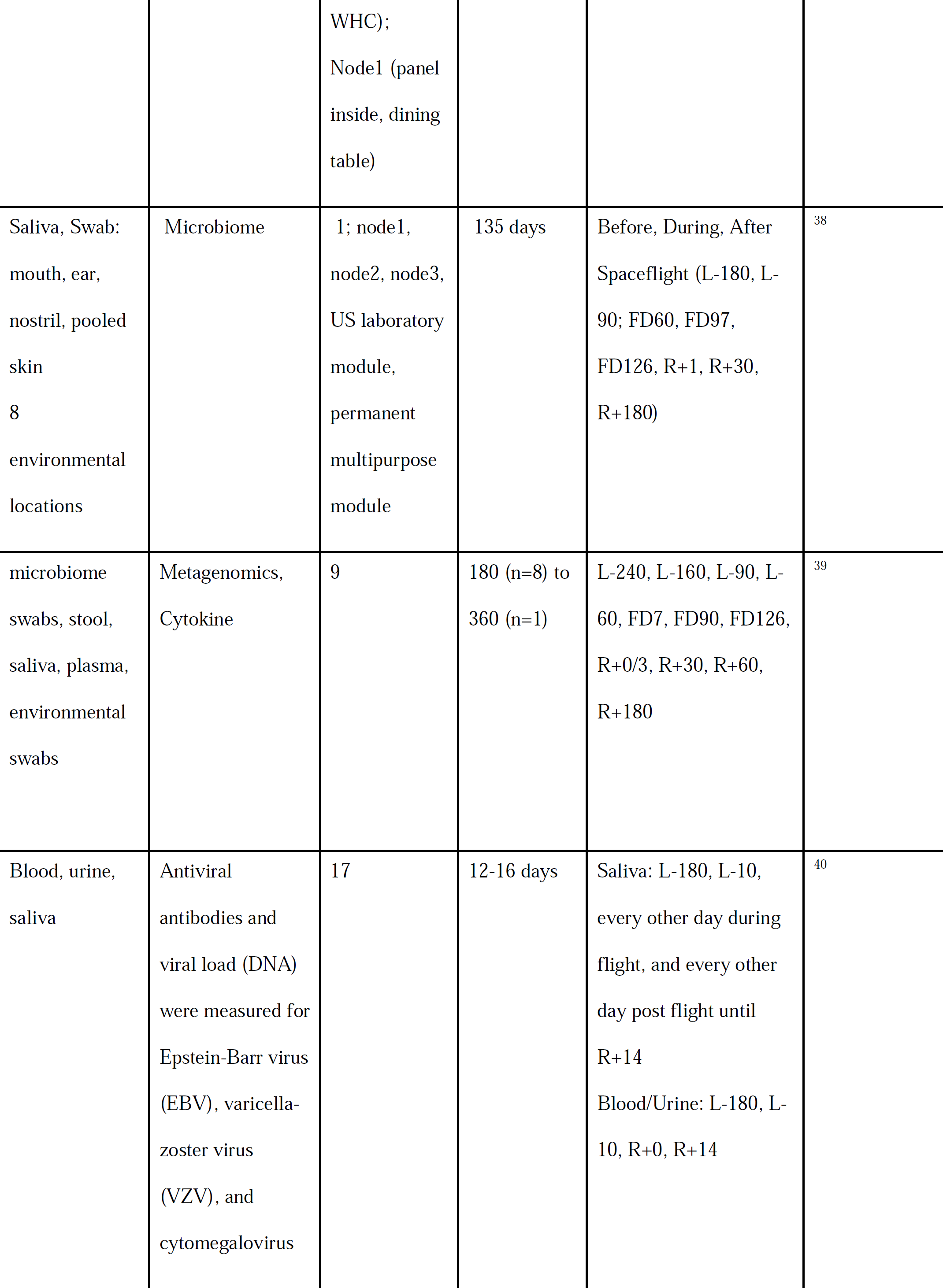
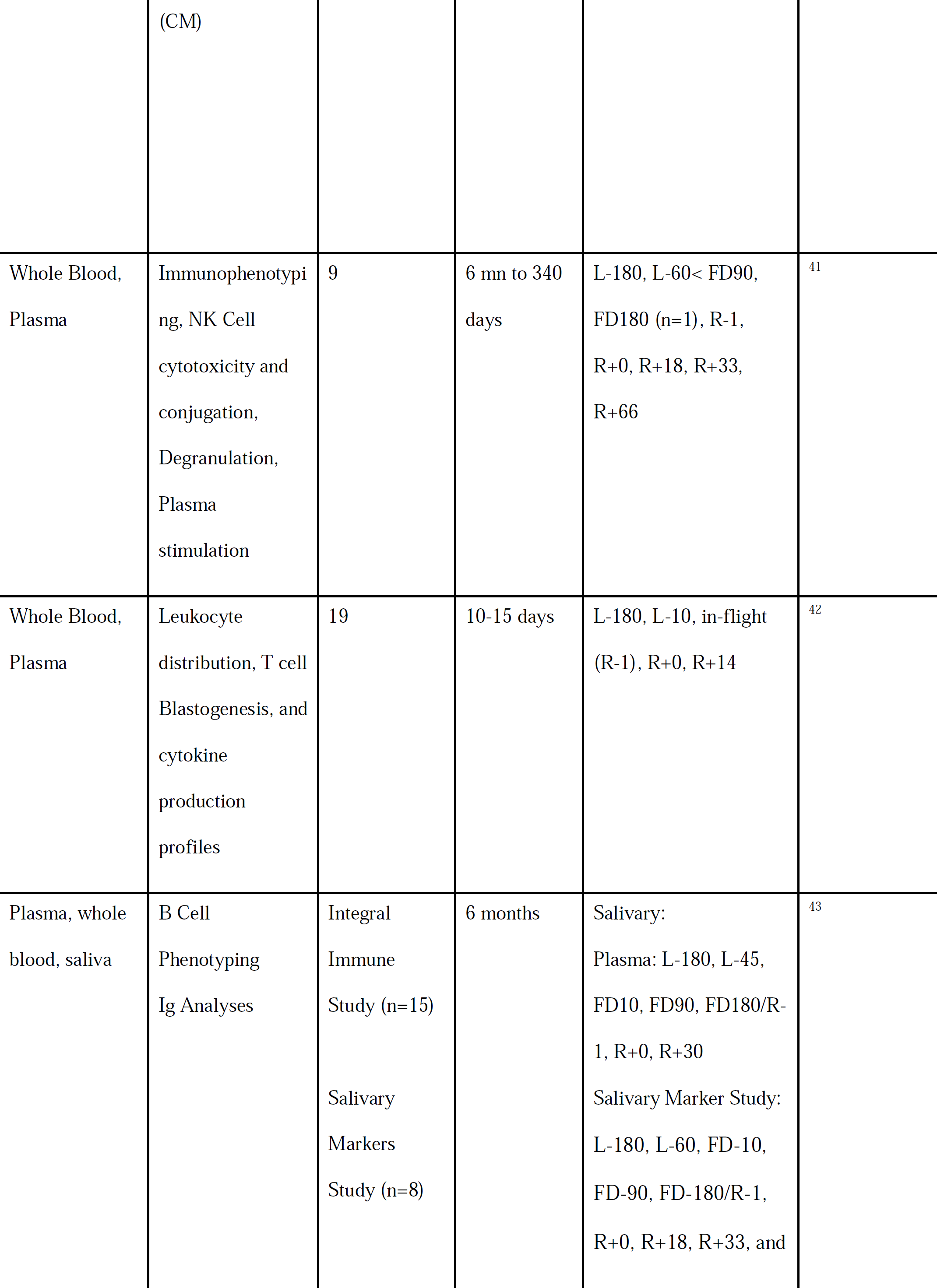
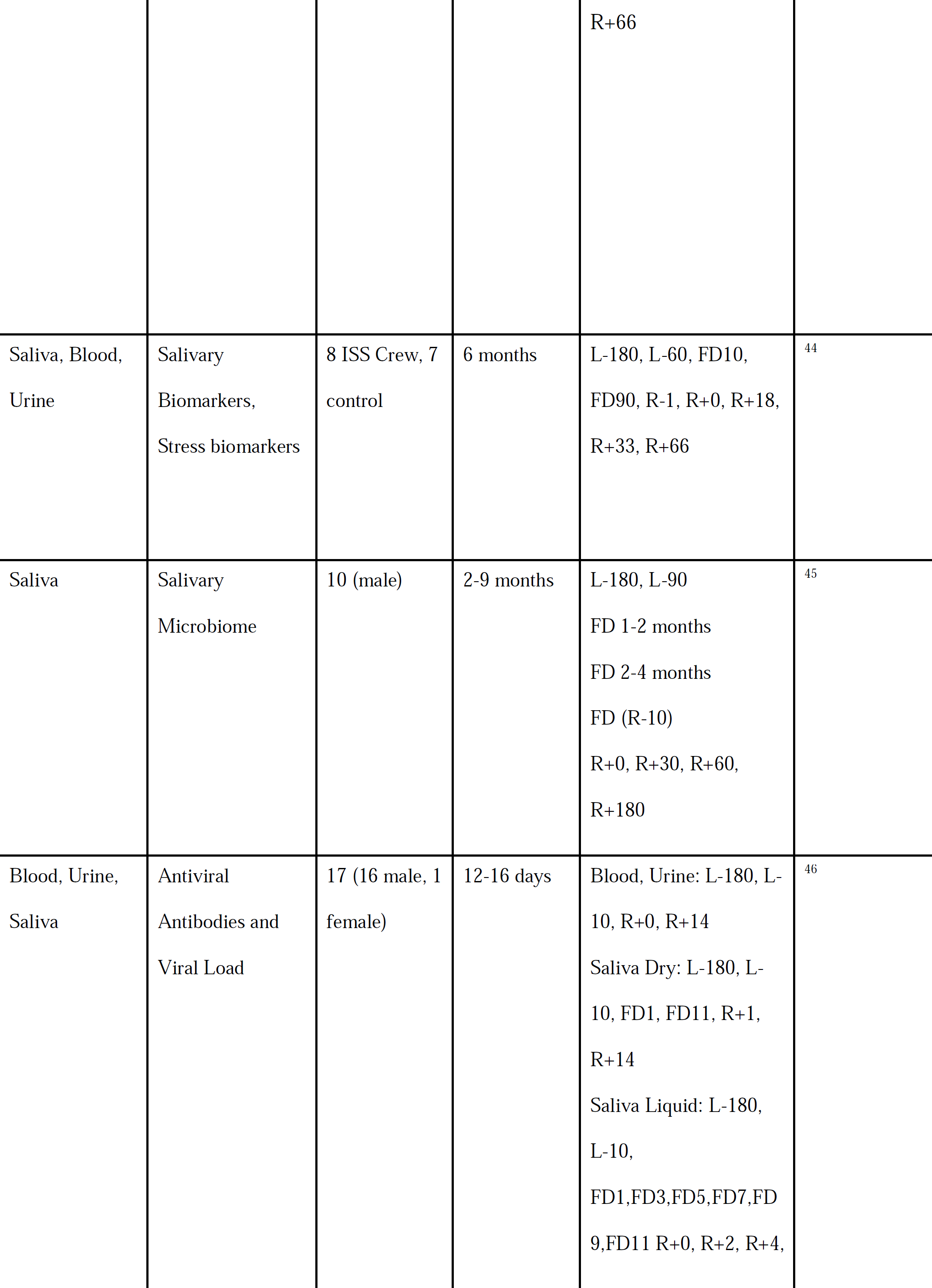
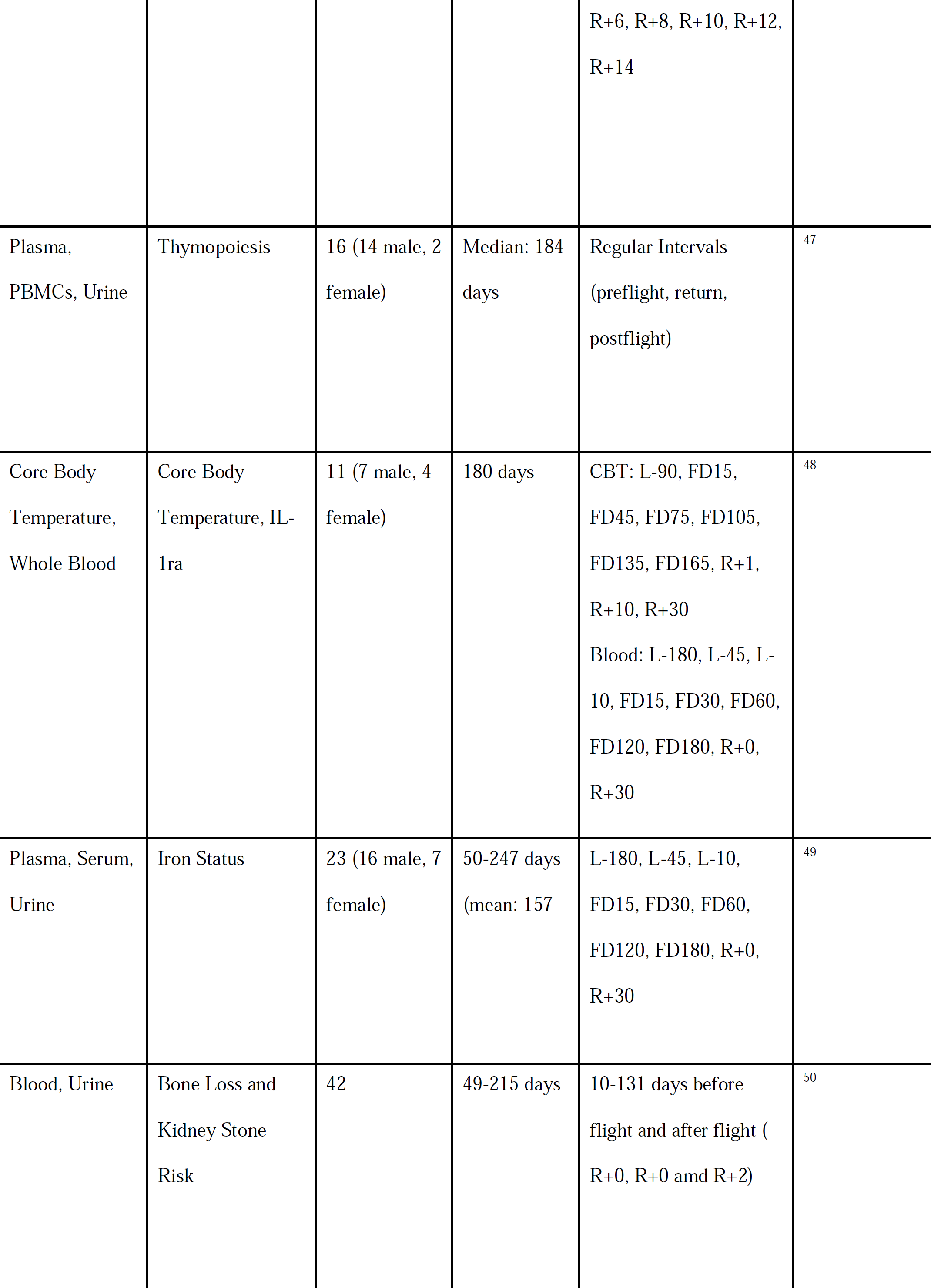
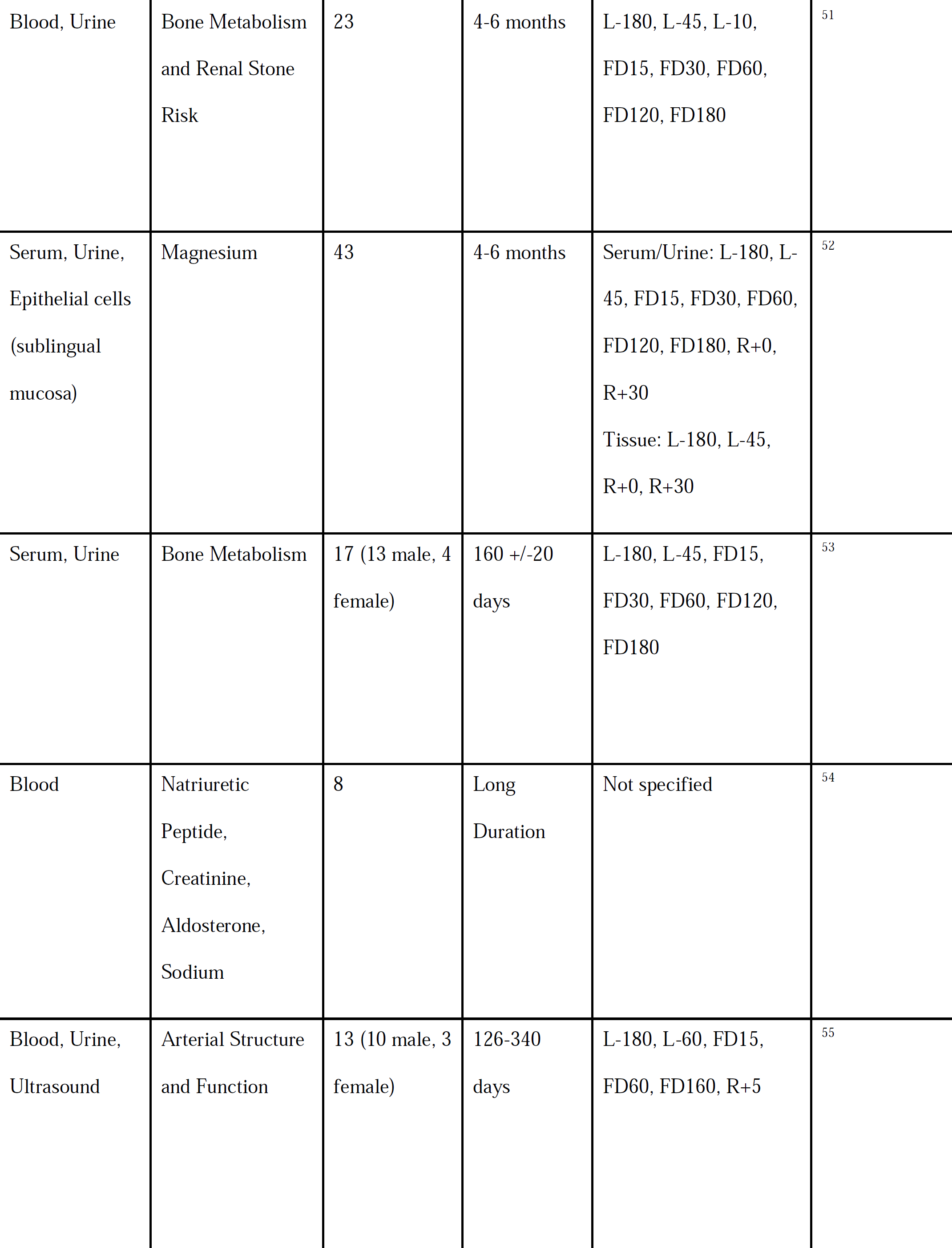
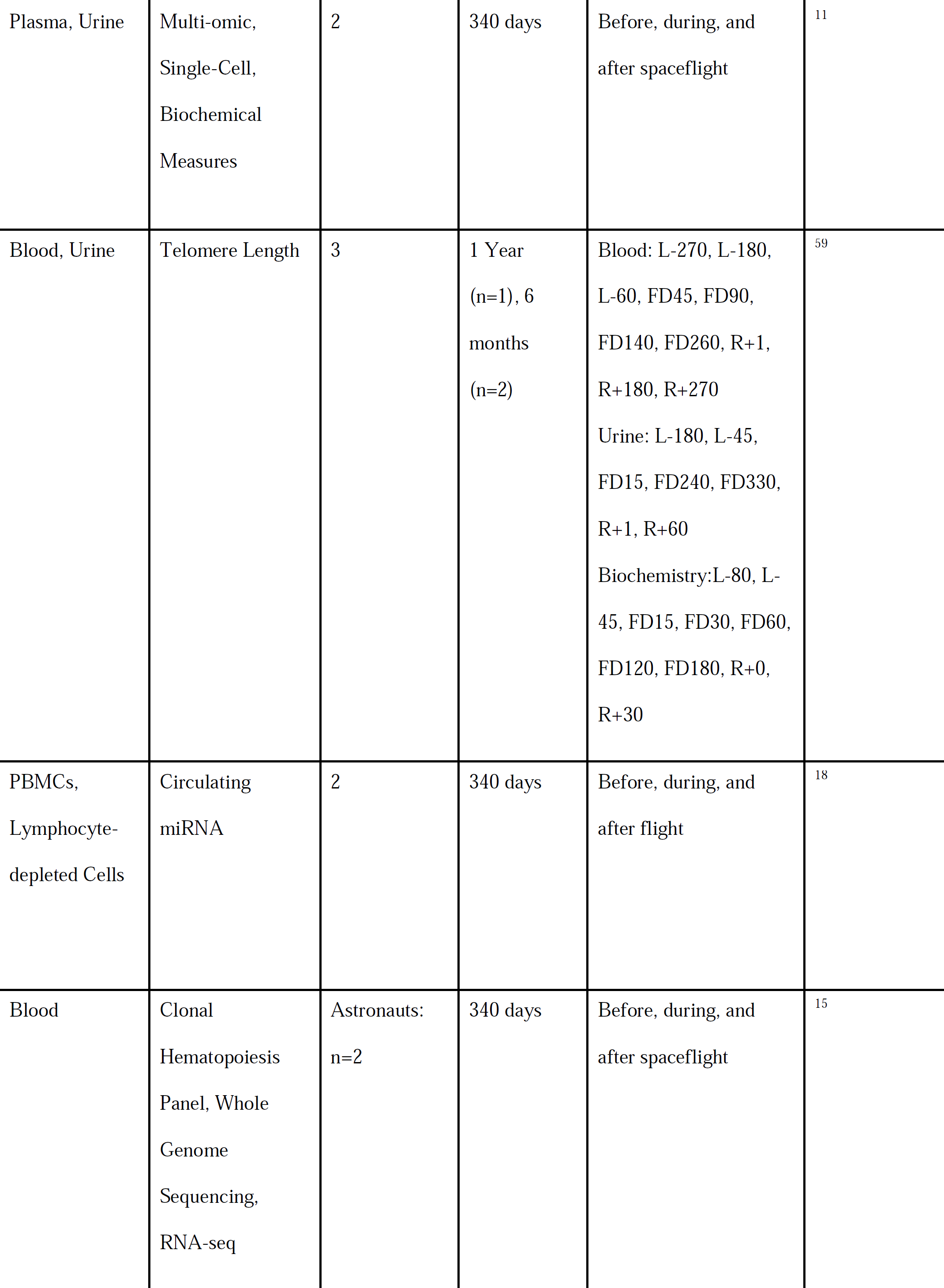
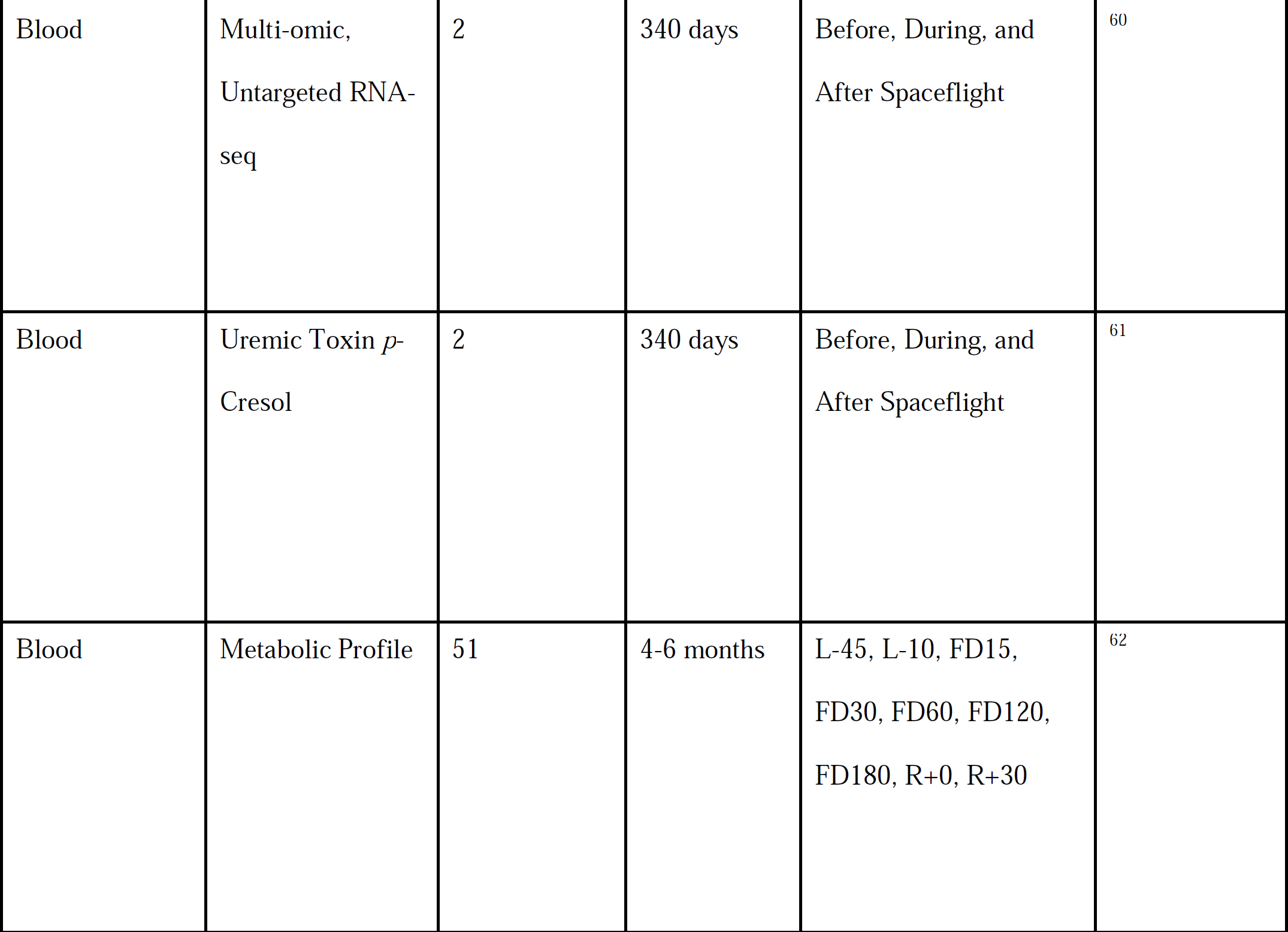
Prior Biospecimen Collections from Astronauts. Listed studies are limited to the past decade.

| Sample(s) | Measure(s) | Number of Subjects (n) | Duration Range (days) | Collection Time points | Study (citation) |
| --- | --- | --- | --- | --- | --- |
| Plasma | mtDNA, Long Non-coding RNA, Exosomes | 3-14 | 5-13 | L-10, R-0, R+3 | <sup>24,25</sup> |
| Plasma, Saliva | Cytokines | 13 | 140-290 | L-180, L-45, L-10, FD15, FD30, FD60, FD120, FD180, R+0, and R+30 | <sup>10</sup> |
| Plasma | Cytokines | 28 | ~180 | L-180, L-45, L-10, FD15,30,60,120,180; R+0, R+30 | <sup>26</sup> |
| Plasma | Proteomics | 13-18 | 169-199 | L-30, R+0, R+7 | <sup>20,27-29</sup> |
| Plasma | sRNAseq<br>(miRNA from<br>sEV) | 14 | 12 (median) | L-10, R+0, R+3 | <sup>30</sup> |
| PBMCs | Peripheral<br>Leukocyte<br>Distribution, T-cell Function,<br>Virus-specific<br>Immunity, and<br>Mitogen-<br>stimulated<br>Cytokine<br>Production<br>profiles | 23 | <60 days<br>(n=2), >100<br>days (n=5), 6<br>months<br>(n=16) | L-180, L-45, FD14, FD<br>2-4 mn, FD6 mn, R+0,<br>R + 30 | <sup>31</sup> |
| plasma,<br>PBMCs | snoRNA<br>Expression Levels | n=5 (plasma),<br>n=6 (PBMCs) | 14 (median) | L-10, R+3 | <sup>32</sup> |
| Whole blood,<br>serum | Hematology | 14 | 167 ± 31<br>days<br>(mean±sd) | L-100, FD5, FD11,<br>FD64, FD157, R+4,<br>R+14, R+41, R+184<<br>R+365 | <sup>13</sup> |
| Whole blood | Transcriptome | 6 | 10-13 | L-10, R+0 (2-3 hour<br>after return) | <sup>33</sup> |
| Whole blood | Hematology | 31 | Up to 180 | L-180, L-45; FD-14,<br>FD60-FD120, FD180,<br>R+0, R+30 | <sup>34</sup> |
| Whole blood,<br>Saliva | Immune Cell<br>Counts, Cortisol | 9 | 162 | L-25, FD90, FD150,<br>R+1, R+7, R+30 | <sup>35</sup> |
| body swabs,<br>saliva | Metagenomics | 4 |  | L-180, L-45; FD-14,<br>FD60-FD120, FD180,<br>R+0, R+30, R+180 | <sup>36</sup> |
| ISS section<br>swab | Metagenomics,<br>Physiological<br>Characterization<br>of Microbes | Locations:<br>Columbus<br>(air, light<br>cover, SSC<br>laptop,<br>handrails,<br>RGSH);<br>Node2<br>(sleeping unit,<br>panel outside,<br>ATU); Cupola<br>(air, surface),<br>Node3<br>(ARED,<br>treadmill, |  | 3 timepoints (session A,<br>B, and C) | <sup>37</sup> |
|  |  | WHC);<br>Node1 (panel<br>inside, dining<br>table) |  |  |  |
| Saliva, Swab:<br>mouth, ear,<br>nostril, pooled<br>skin<br>8<br>environmental<br>locations | Microbiome | 1; node1,<br>node2, node3,<br>US laboratory<br>module,<br>permanent<br>multipurpose<br>module | 135 days | Before, During, After<br>Spaceflight (L-180, L-90; FD60, FD97,<br>FD126, R+1, R+30,<br>R+180) | <sup>38</sup> |
| microbiome<br>swabs, stool,<br>saliva, plasma,<br>environmental<br>swabs | Metagenomics,<br>Cytokine | 9 | 180 (n=8) to<br>360 (n=1) | L-240, L-160, L-90, L-60, FD7, FD90, FD126,<br>R+0/3, R+30, R+60,<br>R+180 | <sup>39</sup> |
| Blood, urine,<br>saliva | Antiviral<br>antibodies and<br>viral load (DNA)<br>were measured for<br>Epstein-Barr virus<br>(EBV), varicella-zoster virus<br>(VZV), and<br>cytomegalovirus | 17 | 12-16 days | Saliva: L-180, L-10,<br>every other day during<br>flight, and every other<br>day post flight until<br>R+14<br>Blood/Urine: L-180, L-10, R+0, R+14 | <sup>40</sup> |
|  | (CM) |  |  |  |  |
| Whole Blood,<br>Plasma | Immunophenotyping, NK Cell cytotoxicity and conjugation, Degranulation, Plasma stimulation | 9 | 6 mn to 340 days | L-180, L-60< FD90, FD180 (n=1), R-1, R+0, R+18, R+33, R+66 | <sup>41</sup> |
| Whole Blood,<br>Plasma | Leukocyte distribution, T cell Blastogenesis, and cytokine production profiles | 19 | 10-15 days | L-180, L-10, in-flight (R-1), R+0, R+14 | <sup>42</sup> |
| Plasma, whole blood, saliva | B Cell Phenotyping<br>Ig Analyses | Integral Immune Study (n=15)<br>Salivary Markers Study (n=8) | 6 months | Salivary:<br>Plasma: L-180, L-45, FD10, FD90, FD180/R-1, R+0, R+30<br>Salivary Marker Study:<br>L-180, L-60, FD-10, FD-90, FD-180/R-1, R+0, R+18, R+33, and | <sup>43</sup> |
|  |  |  |  | R+66 |  |
| Saliva, Blood,<br>Urine | Salivary<br>Biomarkers,<br>Stress biomarkers | 8 ISS Crew, 7<br>control | 6 months | L-180, L-60, FD10,<br>FD90, R-1, R+0, R+18,<br>R+33, R+66 | <sup>44</sup> |
| Saliva | Salivary<br>Microbiome | 10 (male) | 2-9 months | L-180, L-90<br>FD 1-2 months<br>FD 2-4 months<br>FD (R-10)<br>R+0, R+30, R+60,<br>R+180 | <sup>45</sup> |
| Blood, Urine,<br>Saliva | Antiviral<br>Antibodies and<br>Viral Load | 17 (16 male, 1<br>female) | 12-16 days | Blood, Urine: L-180, L-10, R+0, R+14<br>Saliva Dry: L-180, L-10, FD1, FD11, R+1,<br>R+14<br>Saliva Liquid: L-180,<br>L-10,<br>FD1,FD3,FD5,FD7,FD<br>9,FD11 R+0, R+2, R+4, | <sup>46</sup> |
|  |  |  |  | R+6, R+8, R+10, R+12,<br>R+14 |  |
| Plasma,<br>PBMCs, Urine | Thymopoiesis | 16 (14 male, 2<br>female) | Median: 184<br>days | Regular Intervals<br>(preflight, return,<br>postflight) | <sup>47</sup> |
| Core Body<br>Temperature,<br>Whole Blood | Core Body<br>Temperature, IL-1ra | 11 (7 male, 4<br>female) | 180 days | CBT: L-90, FD15,<br>FD45, FD75, FD105,<br>FD135, FD165, R+1,<br>R+10, R+30<br>Blood: L-180, L-45, L-10, FD15, FD30, FD60,<br>FD120, FD180, R+0,<br>R+30 | <sup>48</sup> |
| Plasma, Serum,<br>Urine | Iron Status | 23 (16 male, 7<br>female) | 50-247 days<br>(mean: 157 | L-180, L-45, L-10,<br>FD15, FD30, FD60,<br>FD120, FD180, R+0,<br>R+30 | <sup>49</sup> |
| Blood, Urine | Bone Loss and<br>Kidney Stone<br>Risk | 42 | 49-215 days | 10-131 days before<br>flight and after flight (<br>R+0, R+0 and R+2) | <sup>50</sup> |
| Blood, Urine | Bone Metabolism<br>and Renal Stone<br>Risk | 23 | 4-6 months | L-180, L-45, L-10,<br>FD15, FD30, FD60,<br>FD120, FD180 | <sup>51</sup> |
| Serum, Urine,<br>Epithelial cells<br>(sublingual<br>mucosa) | Magnesium | 43 | 4-6 months | Serum/Urine: L-180, L-45, FD15, FD30, FD60,<br>FD120, FD180, R+0,<br>R+30<br>Tissue: L-180, L-45,<br>R+0, R+30 | <sup>52</sup> |
| Serum, Urine | Bone Metabolism | 17 (13 male, 4<br>female) | 160 +/-20<br>days | L-180, L-45, FD15,<br>FD30, FD60, FD120,<br>FD180 | <sup>53</sup> |
| Blood | Natriuretic<br>Peptide,<br>Creatinine,<br>Aldosterone,<br>Sodium | 8 | Long<br>Duration | Not specified | <sup>54</sup> |
| Blood, Urine,<br>Ultrasound | Arterial Structure<br>and Function | 13 (10 male, 3<br>female) | 126-340<br>days | L-180, L-60, FD15,<br>FD60, FD160, R+5 | <sup>55</sup> |
| Blood, Urine, quantitative CT | Bone Metabolism, Bone Density, Bone Strength | 17 (14 male, 3 female) | 3.5-7 months (mean: 170 days) | Blood/Urine: L-180, L-45, FD15, FD30, FD60, FD120, FD180, R+0 | <sup>56</sup> |
| Stool, Saliva, Skin, Urine, Blood, Plasma, PBMCs | Metabolomics, Proteomics, Cognition, Microbiome, Telomeres, Epigenomics, Biochemical Profile, Gene Expression, Integrative Omics, Immunome | 2 | 1-Year (340 days) | Before, during, and after spaceflight | <sup>17</sup> |
| Blood, Urine | Multi-omics | 59 | 4-6 months | L-180, L-45, FD15, FD30, FD60, FD120, FD180, R+0, R+30 | <sup>57</sup> |
| Plasma | Cell-free DNA, Exosome | 2 | 340 days | Before, during, and after spaceflight (12 timepoints from twin on earth and 11 from twin in space) | <sup>58</sup> |
| Plasma, Urine | Multi-omic,<br>Single-Cell,<br>Biochemical<br>Measures | 2 | 340 days | Before, during, and<br>after spaceflight | <sup>11</sup> |
| Blood, Urine | Telomere Length | 3 | 1 Year<br>(n=1), 6<br>months<br>(n=2) | Blood: L-270, L-180,<br>L-60, FD45, FD90,<br>FD140, FD260, R+1,<br>R+180, R+270<br>Urine: L-180, L-45,<br>FD15, FD240, FD330,<br>R+1, R+60<br>Biochemistry:L-80, L-45, FD15, FD30, FD60,<br>FD120, FD180, R+0,<br>R+30 | <sup>59</sup> |
| PBMCs,<br>Lymphocyte-<br>depleted Cells | Circulating<br>miRNA | 2 | 340 days | Before, during, and<br>after flight | <sup>18</sup> |
| Blood | Clonal<br>Hematopoiesis<br>Panel, Whole<br>Genome<br>Sequencing,<br>RNA-seq | Astronauts:<br>n=2 | 340 days | Before, during, and<br>after spaceflight | <sup>15</sup> |
| Blood | Multi-omic,<br>Untargeted RNA-seq | 2 | 340 days | Before, During, and<br>After Spaceflight | <sup>60</sup> |
| Blood | Uremic Toxin *p*-<br>Cresol | 2 | 340 days | Before, During, and<br>After Spaceflight | <sup>61</sup> |
| Blood | Metabolic Profile | 51 | 4-6 months | L-45, L-10, FD15,<br>FD30, FD60, FD120,<br>FD180, R+0, R+30 | <sup>62</sup> |

For the Inspiration4 mission, sample collection spanned three time points pre-launch (L-92, L-44, L-3 days), three time points during flight (Flight Day 1 (FD1), FD2, FD3), and four time points post-return (R+1, R+45, R+82, R+194 days). Venous blood, urine, stool, and skin biopsies were collected during ground timepoints only, while capillary DBSs, saliva, and skin swabs were collected both on the ground and during flight (**Fig 1b**). Environmental swabs of the Dragon capsule were collected pre-flight in the crew training capsule and during flight in the spacecraft launched from Cape Canaveral (**Fig 1b**).

Samples were collected across a variety of locations based on the crew’s training and travel schedule. L-92 and L-44 were collected in Hawthorne, CA at SpaceX Headquarters, L-3 and R+1 were collected at Cape Canaveral, FL at a facility near the launch-site. FD1, FD2, and FD3 were collected inside the Dragon capsule while in orbit. R+45 was collected at the crew members’ individual locations (which spanned the US States NY, NJ, TN, and WA), R+82 was collected at Weill Cornell Medicine, NY and R+194 was collected at Baylor College of Medicine, TX (**Fig 1c**).

### Blood Collection and Derivatives

Blood was collected using a combination of venipuncture tubes to collect venous blood and contact-activated lancets to collect capillary blood from the fingertip. Each crew member provided blood samples, collected into one blood RNA tube (bRNA), four K2 EDTA tubes, two cell preparation tubes (CPTs), one cell-free DNA tube (cfDNA BCT), one serum separator tube (SST), and one dried blood spot (DBS) card per time point. From these tubes, whole blood, plasma, PBMCs, serum, and cell pellet samples were collected (**Table 2**). Sample yields are reported below. Samples were aliquoted for long-term storage and biobanking (**Table 3**).

**Table 2:** Blood Derivative Allocations. Samples types collected, their tube type of origin, and assay allocation. Samples collected in excess were biobanked to enable additional experiments as new assays are developed.

| Sample Type | Tube Source | Assay Allocation(s) |
| --- | --- | --- |
| Whole Blood | bRNA | Total RNA Extraction |
| Plasma | CPT | Proteomics, Metabolomics; Biobanking |
| PBMCs | CPT | Biobanking |
| Red Blood Cell Pellet | CPT | gDNA; Biobanking |
| Serum | SST | Immune and Cardiovascular Disease Panel, Metabolic Panel; Biobanking |
| Red Blood Cell Pellet | SST | gDNA; Biobanking |
| Plasma | cfDNA BCT | cfDNA; Biobanking |
| Red Blood Cell Pellet | cfDNA BCT | gDNA; Biobanking |
| PBMCs | K2 EDTA | Single-Cell Multiome GEX+ATAC and BCR/TCR Immune Repertoire Profiling |
| Plasma | K2 EDTA | EVPs |
| Whole Blood | K2 EDTA | Complete Blood Count |

**Table 3:**
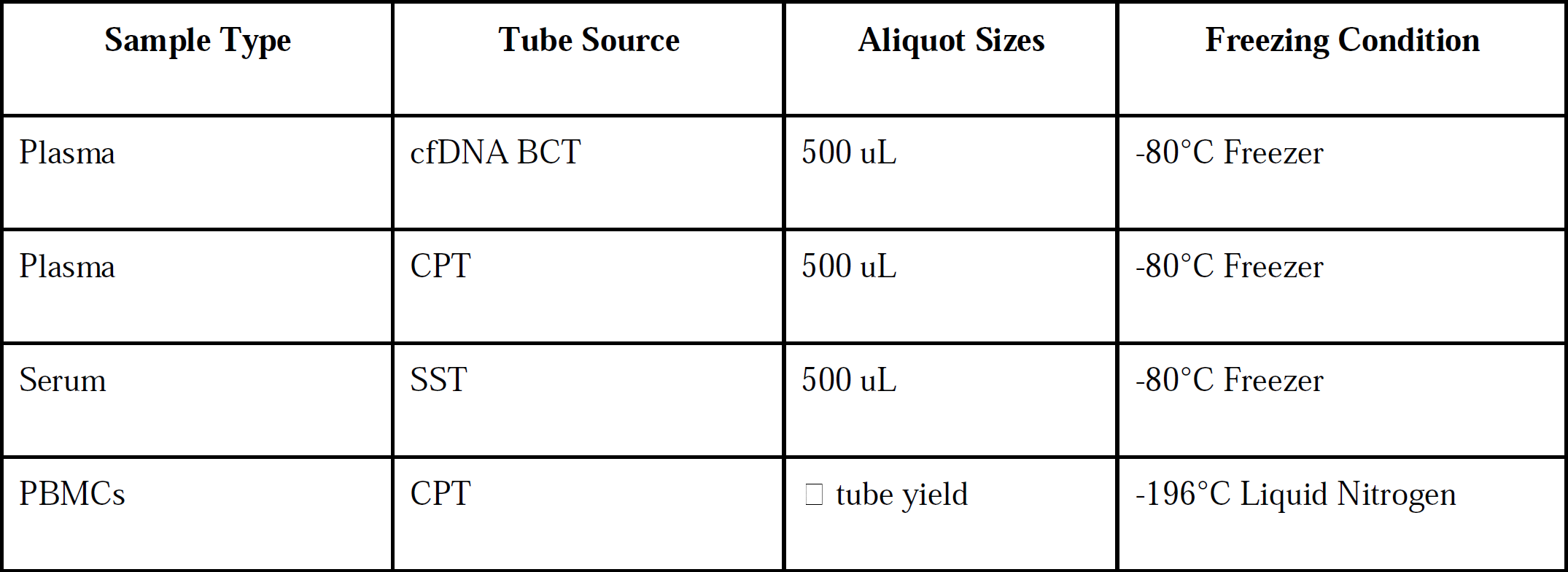
Blood Derivative Aliquot Parameters. Plasma, serum, and PBMCs aliquots were created for downstream assays that only require a portion of the total sample collected in order to minimize freeze-thaw cycles.

bRNA tubes were collected in order to isolate total RNA using the PAXgene blood RNA kit (**Fig 2a**). Yield ranged from 3.04-14.04 µg/tube of total RNA across all samples and the RNA integrity number (RIN) ranged from 3.2-8.5 (mean: 6.95) (**Fig 2b**). RNA was stored at −80°C after extraction. The collection of total RNA enables a variety of downstream RNA profiling methods. It will allow comparative studies to prior RNA-sequencing performed on astronauts, particularly snoRNA & lncRNA biomarkers analyzed from Space Shuttle era blood^25, 32^, mRNA & miRNA measured during the NASA Twin Study^17, 18^, and whole blood RNA arrays from the ISS^33^. Additionally, RNA yields are more than sufficient to perform direct-RNA sequencing using Oxford Nanopore Technologies (ONT) platforms, which require 500 ng of total RNA per library (Manufacturer’s protocol, ONT kit SQK-RNA002). This enables the study of RNA modification changes during spaceflight to create epitranscriptomic profiles for the first time in astronauts.

**Figure 2:**
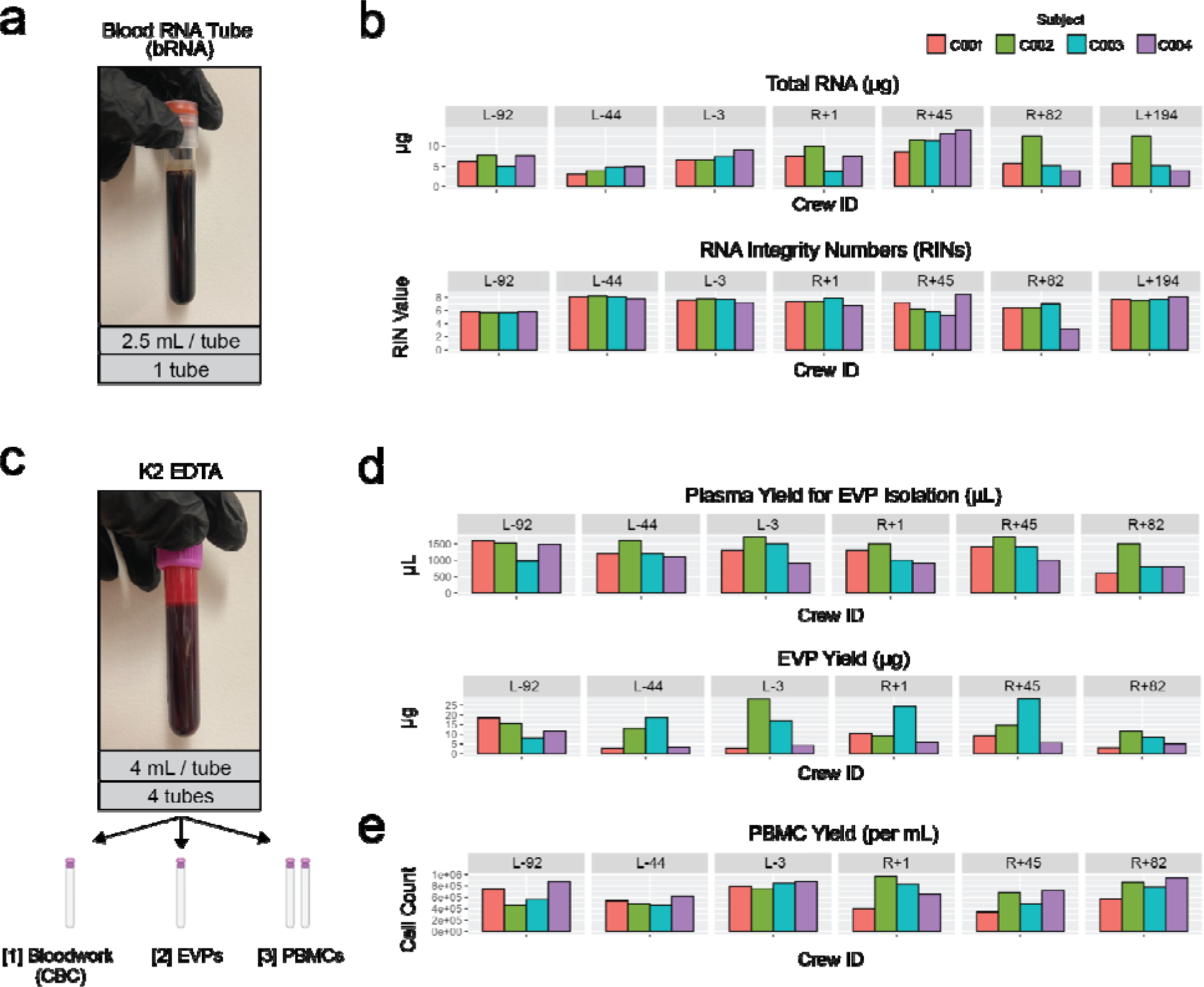
bRNA and K2 EDTA Tubes. **(a)** One 2.5mL bRNA tube was collected per crew member at each ground timepoint. **(b)** bRNA tube total RNA yields per sample (μg) and RINs. **(c)** Four K2 EDTA tubes were collected per member at each ground timepoint. One tube was used for a CBC, one tube was used to isolate EVPs, and two tubes were used for isolation of PBMCs. **(d)** Plasma and EVP yields from the “[2] EVPS” tube on figure 2c. **(e)** PBMC yields per mL from the “[3] PBMCs” tubes on figure 2c.

Four K2 EDTA tubes were drawn at each timepoint from each crew member (**Fig 2c**). One K2 EDTA tube was submitted to Quest Diagnostics to perform a complete blood count (CBC, Quest Test Code: 6399). One tube was used to isolate extracellular vesicles and particles (EVPs) for proteomic quantification (**Fig 3a**). Total EVP quantities varied from 2.71-28.27 ug (**Fig 2d**). Two K2 EDTA tubes were used to isolate PBMCs for single-cell sequencing (10X Chromium Single Cell Multiome ATAC + Gene Expression and Chromium Single Cell Immune Profiling workflows). After collection, a Ficoll separation was performed to isolate PBMCs, which ranged from 340,000-975,000 cells per mL of blood (**Fig 2e**). One prior single-cell gene expression experiment, NASA Twin study, was performed on astronauts, which found immune cell population specific gene expression changes and a correlation with microRNA signatures^11, 18^.

**Figure 3:**
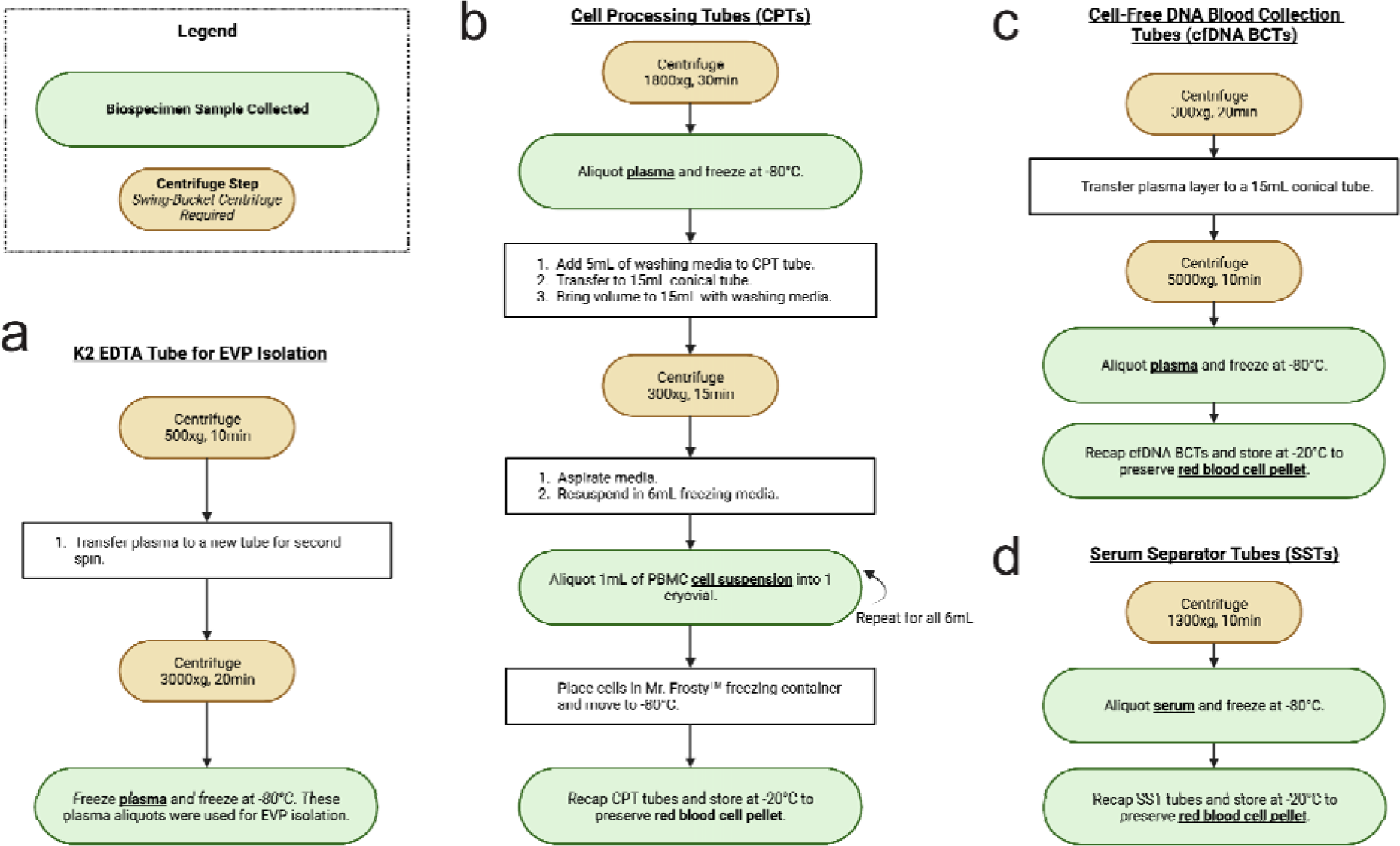
Tube Processing Steps. Centrifuge (brown circles) and aliquoting (white and green boxes and circles) protocols for **(a)** K2 EDTA tubes designated for EVP isolation **(b)** CPTs **(c)** cfDNA BCTs and **(d)** SSTs.

Additional PBMCs, plasma, and serum were collected from CPTs (**Fig 4a**), cfDNA BCTs (**Fig 4d**), SSTs (**Fig 4c**), as well as red blood cell pellets. CPTs were spun and aliquoted according to the manufacturer’s instructions (**Fig 3b**). Plasma volume per tube ranged from 3000-14,000 uL per tube (**Fig 4d**). There were a few instances were CPT tubes shattered in the centrifuge and plasma could not be salvaged. Plasma can be used to validate or refute previous studies, including cytokine panel^10, 26^, exosomal RNA-seq^25, 32^, extracellular vesicle microRNA^30^, and proteomic^20, 27–29^ results. PBMCs were also collected, aliquoted into 6 cryovials per CPT, and stored in liquid nitrogen after slowly cooled in a Mr. Frosty to −80°C. These can be used to follow-up on previous studies on adaptive immunity, cell function, and immune dysregulation^8, 31, 41–43^. The remaining red blood cell pellet mixtures from below the gel plug in each CPT Tube were stored at −20°C.

**Figure 4:**
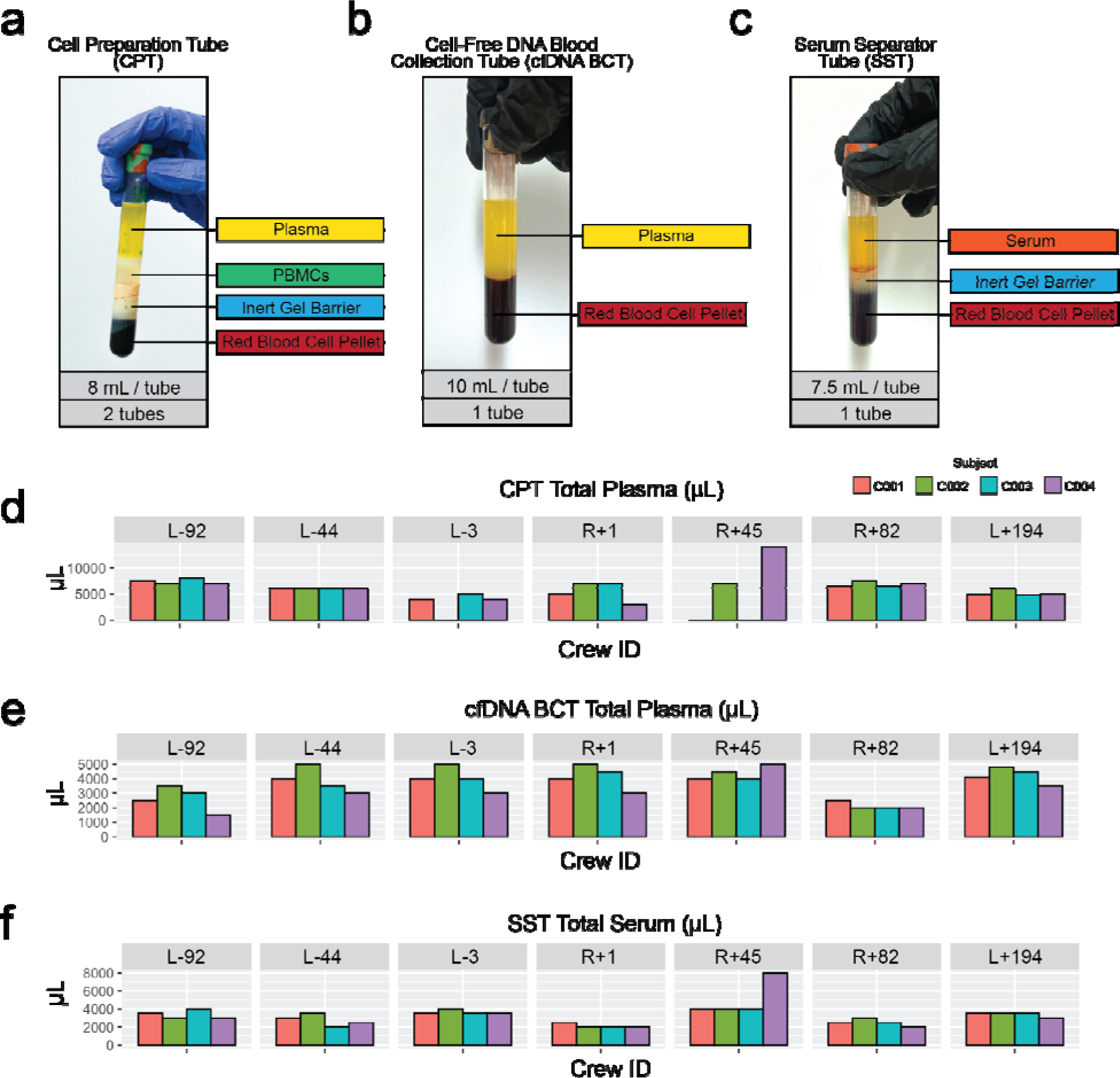
CPT, cfDNA BCT, and SST Yields. **(a)** A spun CPT yields plasma, PBMCs, and a red blood cell pellet. PBMC from each tube were divided into 6 cryovials and viably frozen. Plasma was aliquoted and the pellet was frozen at −20C. **(b)** A spun cfDNA BCT yields plasma and a red blood cell pellet. Plasma was purified with an additional spin (see Fig 4a) then aliquoted. The pellet was frozen at −20C. **(c)** A spun SST yields serum and a red blood cell pellet. Serum was aliquoted and the pellet was frozen at −20C. **(d)** CPT plasma volumes per timepoint are reported. **(e)** cfDNA BCT plasma volumes per timepoint. **(f)** SST serum volumes per timepoint. An extra tube was drawn for C004 at R+45, resulting in a higher serum yield.

cfDNA BCT tubes were collected to isolate high-quality cfDNA from plasma. cfDNA BCTs were spun and aliquoted according to the manufacturer’s instructions (**Fig 3c**). The remaining cell pellet mixture was frozen at −20°C. Plasma volume per timepoint ranged from 1500-5000 uL (**Fig 4e**). 500 uL aliquots were frozen at −80°C. cfDNA extracted from these tubes can be analyzed for fragment length, mitochondrial or nuclear origin, and cell type or tissue of origin^24, 58^.

The SST was spun and aliquoted according to the manufacturer’s instructions (**Fig 3d**). Serum volume ranged from 2000-8000 uL per timepoint (**Fig 4f**). Similar to plasma, serum can be allocated for cytokine analysis and can also be used to perform comprehensive metabolic panels, including one we used at Quest (CMP, Quest Test Code: 10231) for metrics on alkaline phosphatase, calcium, glucose, potassium, and sodium, among other metabolic markers. The remaining cell pellet mixture from each SST tube was stored at −20°C.

In addition to venous blood, capillary blood was collected onto a DBS card using a contact-activated lancet pressed against the fingertip (**Fig 5a**). Capillary blood was collected onto a dried blood spot (DBS) card to preserve nucleic acids and proteins. The amount of capillary blood collected across timepoints varied (**Fig 5b, 5c**) according to how much blood could be collected before the puncture wound closed.

**Figure 5:**
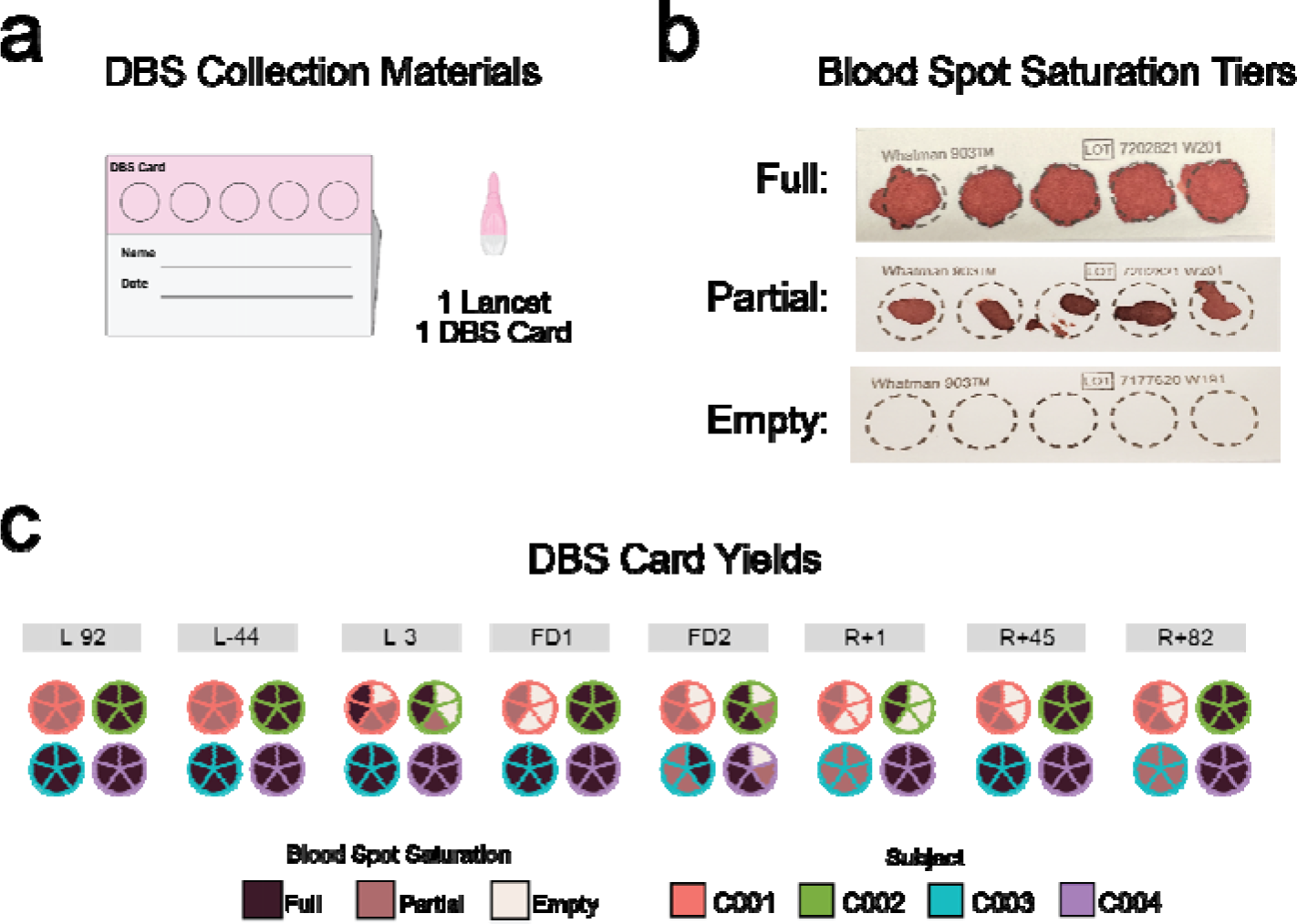
DBS Collection Yields. **(a)** Dried blood spot cards were collected preflight, during flight, and postflight. There were five spots for blood collection per card. **(b)** Blood collections varied in saturation level across blood spots and timepoints. These were classified as “full”, “partial”, and occasionally “empty”. **(c)** DBS card yields per blood spot, per timepoint, and per crew member.

### Saliva Collection

Saliva was collected at the L-92, L-44, L-3, FD1, FD2, FD3, R+1, R+45, and R+82 timepoints using two methods. First, saliva was collected using the OMNIgene Oral Kit, which preserves nucleic acids (**Fig 6a**) during the ground timepoints. From these samples, DNA, RNA, and protein were extracted. DNA yield ranged from 28.1 to 3,187.8 ng, RNA yield from 396.0 to 3544.2 ng (less the two samples had concentrations too low for measurement), and protein concentration from 92.97 - 93.15 ng.

**Figure 6:**
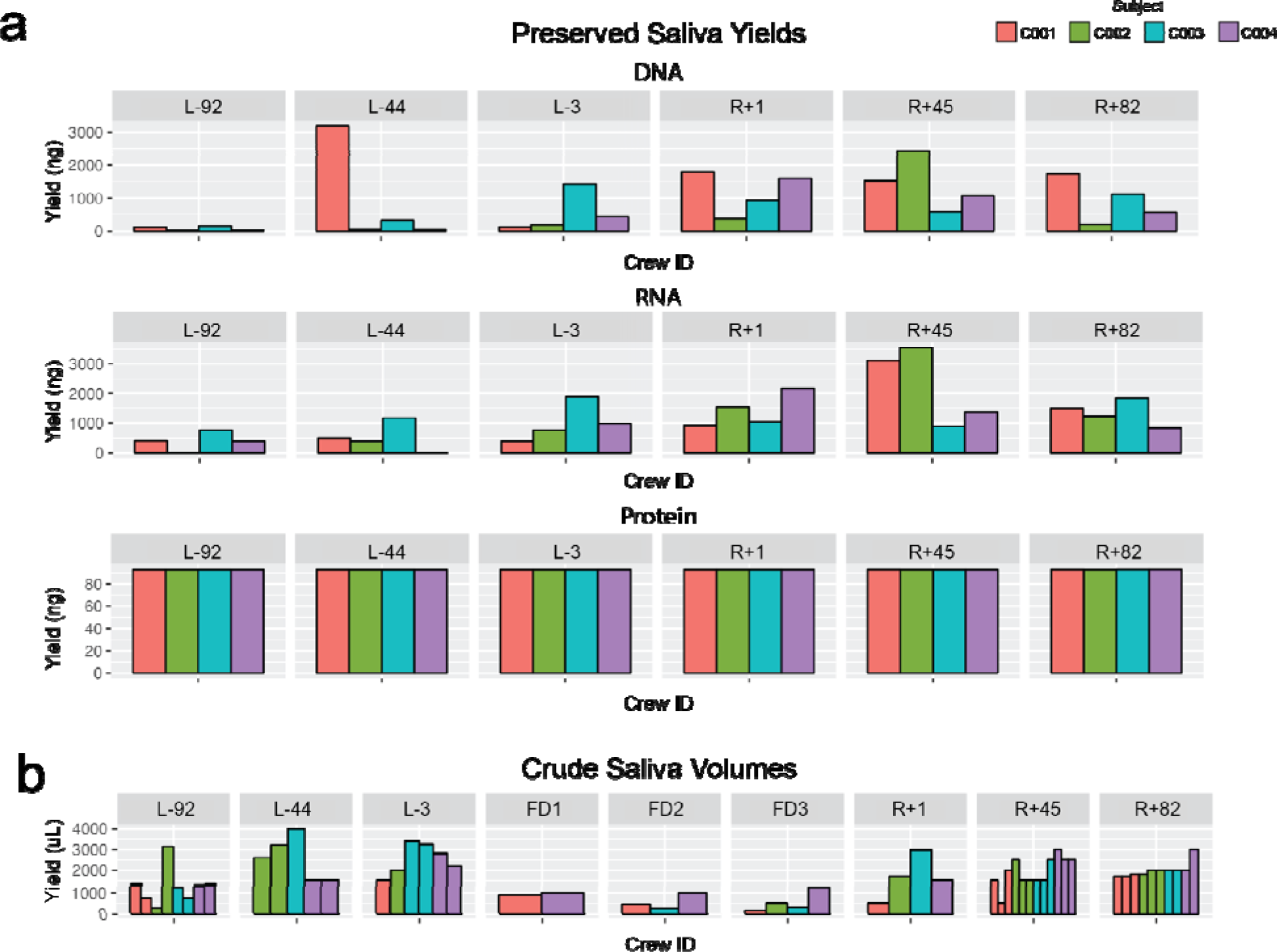
Saliva, Urine, and Stool Sample Collections. **(a)** DNA, RNA, and protein yields from the OMNIgene Oral kits. **(b)** Volume of crude saliva collected per timepoint.

Second, crude saliva (i.e. saliva with no preservative added) was collected into a 5mL DNase/RNase-free screw top tube during the ground and flight timepoints. Saliva volume varied from 150 - 4,000 uL per tube (**Fig 6b**). Crude saliva was also collected during flight (FD2 and FD3), in addition to the ground timepoints.

Saliva collections have been conducted throughout spaceflight studies for assessing the immune state, particularly in the context of viral reactivation. Previously identified viruses that reactivate during spaceflight include Epstein–Barr, varicella-zoster, and cytomegalovirus ^46^. Responses to reactivation of these viruses can be asymptomatic, debilitating, or even life-threatening, thus assessing these adaptations is beneficial in understanding viral spaceflight activity as well as crew health. In addition to viral nucleic acid quantification, numerous biochemical assays can also be performed, including measurements of C-reactive protein (CRP), cortisol, dehydroepiandrosterone (DHEA), and cytokines, among others ^10, 35, 44, 46^.

### Urine Collection

Urine was collected in sterile specimen cups at the L-92, L-44, L-3, R+1, R+45, and R+82 timepoints. Specimen cups were collected 1-2 times per day. For preservation, urine was aliquoted and stored at −80°C. Half the urine had Zymo Urine Conditioning Buffer (UCB) added before freezing, to preserve nucleic acids. Samples yielded 23 - 155.5 mL of crude urine and 21 - 112 mL of UCB urine per specimen cup (**Fig 7a**). Urine was split into 1 mL - 15 mL aliquots before freezing at −80°C.

**Figure 7:**
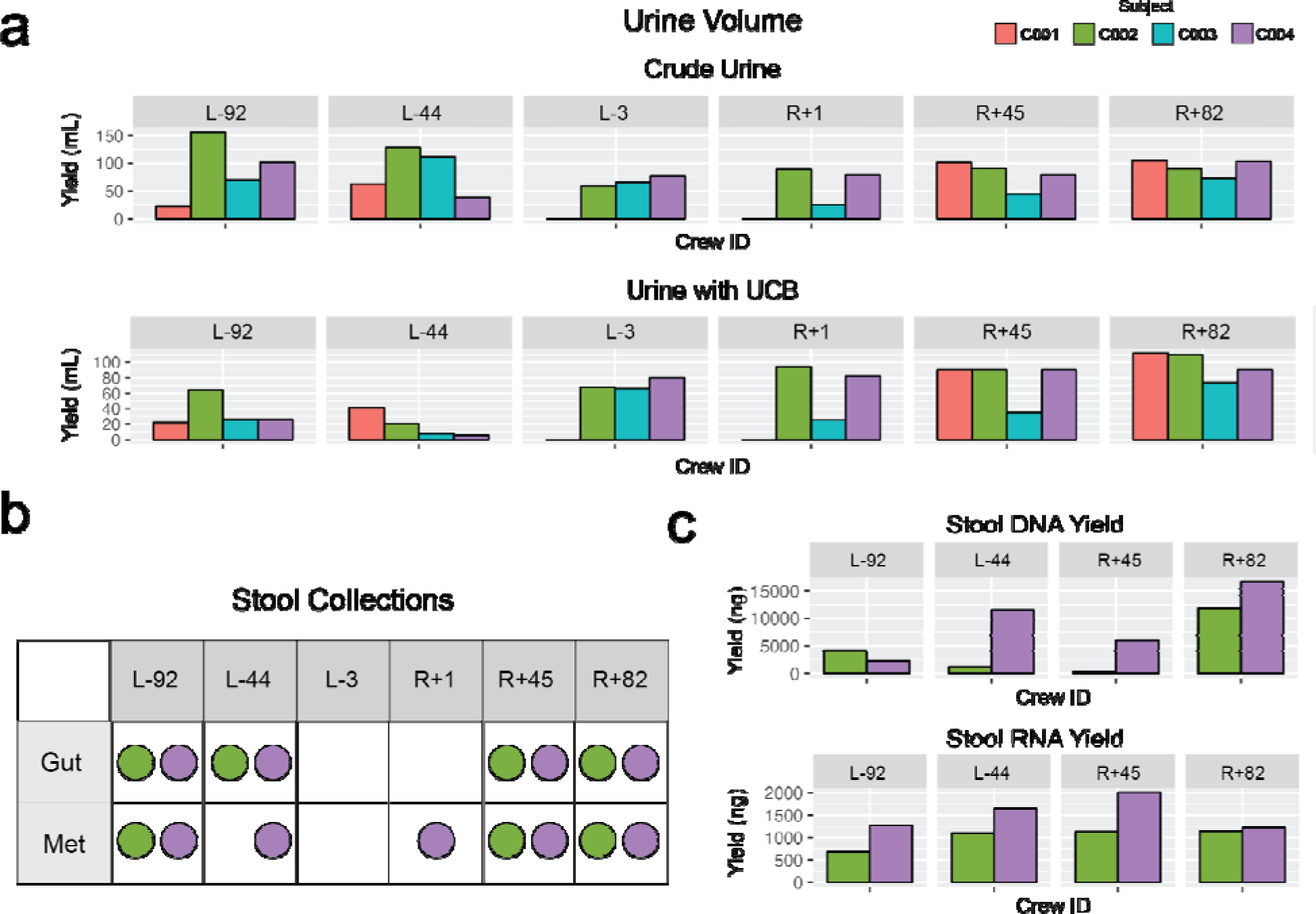
Urine and Stool Sample Collections. **(a)** Urine volumes per timepoint. Volumes are reported for both crude urine and urine preserved with Zymo urine conditioning buffer (UCB). **(b)** Timepoints that stool tubes were collected. “Gut” tubes are OMNIgene•GUT tubes for microbiome preservation. “Met” tubes are OMNImet•GUT tubes for metabolome preservation. **(c)** Stool “Gut” tube DNA and RNA extraction quantities.

A wide variety of assays can be performed on urine samples. Previous studies have included viral reactivation^40, 44, 46^, urinary cortisol^47, 55^, iron and magnesium measurements^49, 52^, bone status^50, 51, 53, 56^, kidney stones^50, 51^, proteomics^11^, telomere measurements^59^, and various biomarkers and metabolites^17, 55^.

### Stool Collection

Stool was collected at the L-92, L-44, R+1, R+45, and R+82 timepoints. Stool samples were stored into two collection containers at each timepoint, one DNA Genotek OMNIgene Gut (OMR-200) kit with a preservative for metagenomics and another (ME-200) with a preservative for metabolomics (**Fig 7b**). Stool was the least consistent sample collected due to the limited windows available for sampling during collection timeframes. DNA and RNA were extracted from aliquots of the OMNIgene Gut (OMR-200) tubes for downstream microbiome analysis. DNA yield ranged from 358.5 - 16,660 ng, RNA from 690 - 2010 ng (**Fig 7c**). Large variations in yield are attributable to variable stool mass collected between kits.

Stool samples enable various biochemical, immune, and microbiome changes studies. Previous metagenomic assays have found that shannon alpha diversity and richness during long duration missions to the ISS ^39^.

### Skin Swabs

Body swabs were collected at all timepoints. Samples were collected by swabbing the body region of interest for 30 seconds, then placing the swab in a sterile 2D matrix tube (Thermo Scientific #3710) with Zymo DNA/RNA shield preservative. For the first two swab locations, the oral and nasal cavity, the swab was placed directly on the body after removal from its sterile packaging (dry-swab method; **Fig 8a**). For the remaining body locations, the swab was briefly dipped in nuclease-free, DNA/RNA-free water before proceeding (wet-swab method). Eight distinct sites were swabbed with the wet-swab method: post-auricular, axillary vault, volar forearm, occiput, umbilicus, gluteal crease, glabella, and the toe-web space (**Fig 8b**). The astronaut microbiome has previously been studied in the forehead, forearm, nasal, armpit, navel, postauricular, and tongue body locations, and changes have been documented during flight. Changes in alpha diversity and beta diversity were documented, as well as shifts in microbial genera^39^. However, the impact of these changes on skin health and immunological health are not well understood.

**Figure 8:**
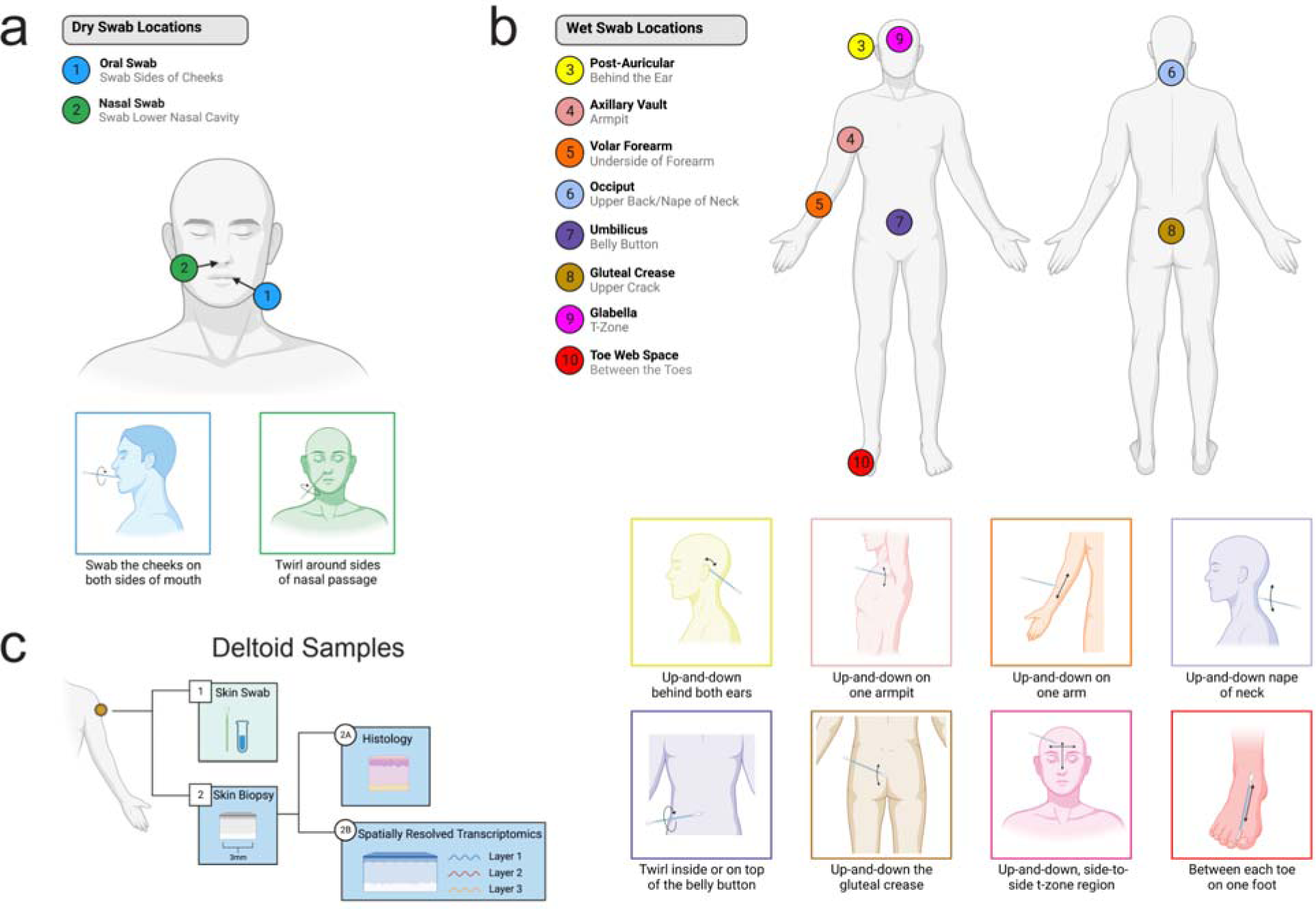
Skin Collection Locations and Sample Types. **(a)** Dry swabs were collected from two body locations. **(b)** Wet swabs were collected from eight body locations. **(c)** Swabs were collected from the deltoid region. Immediately after, 3- or 4-mm skin biopsies were collected from the same area and divided for histology and spatially resolved transcriptomics.

Acquiring extensive swab samples from the crew skin allows for characterization of the habitat environment, crew skin microbiome adaptations, and interactions with potential human health adaptations resulting from spaceflight exposure. This is very relevant for crew health, considering astronauts become more susceptible to infections during spaceflight missions^63^, with the relationship between microbe-host interactions from spaceflight exposure, which may be a causative factor of astronauts immune dysfunction, which is still not well understood.

### Skin Biopsies

A skin biopsy on the deltoid was obtained from the L-44 and R+1 timepoint. Biopsies were also collected in advance of a flight to ensure the biopsy site is fully healed before the flight so there is no risk of complication.The wet-swab method was used to collect the skin microbiome before the skin biopsy. The skin biopsies were three millimeters in diameter and were collected for histology and spatially resolved transcriptomics (SRT) (**Fig 8c**). One-third of the sample was stored in formalin and kept at room temperature to perform histology. The remaining two-thirds of the sample was stored in a cryovial and placed at −80°C for SRT (**Fig 8c**). This is the first sample collected from astronauts for spatially resolved transcriptomics. The skin is of high interest due to the inflammation-related cytokine markers such as IL-12p40, IL-10, IL-17A, and IL-18^10, 17^ and skin rash’s status as the most frequent clinical symptom reported during spaceflight^64^.

### Environmental Swabs and HEPA Filter

Environmental swabs were collected in flight during the F1 and F2 timepoint. Additionally, environmental swabs were collected from the flight simulation capsule at SpaceX headquarters after days of crew training during the L-92 and L-44 timepoints. Environmental swabs were collected using the wet-swab method. Ten environmental swabs were collected per time point at the following locations in the capsule: an ambient air/control swab, the execute button, the viewing dome, the side hatch mobility aid, the lid of the waste locker, the head section of one of the seats, the commode panel, the right and left sides of the control screen, and the g-meter button (**Fig 9a-d**). Additionally, the spacecraft’s high-efficiency particulate absorbing (HEPA) filter was acquired post-flight (**Fig 10a**). This filter was cut into 127 rectangular pieces (1.2” × 1.6” × 4”) and stored at −20°C (**Fig 10b**, **Fig 10c**).

**Figure 9:**
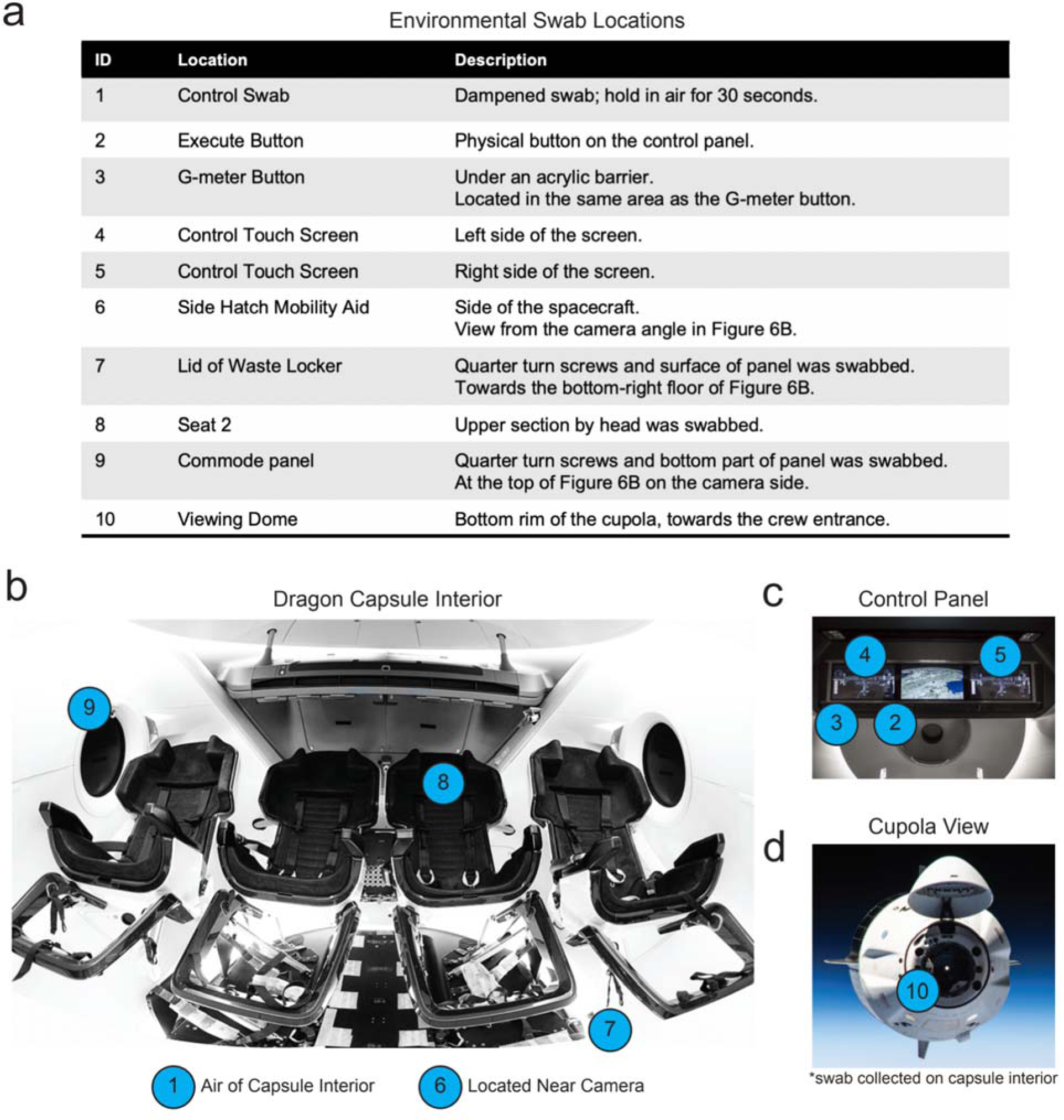
Capsule Swab Locations. **(a)** Swab locations, descriptions, and label IDs. **(b)** Interior view of the SpaceX Dragon capsule. **(c)** View of the control panel located above the middle seats in the Dragon capsule. **(d)** View of the cupola (viewing dome) region from the outside. The rim of the dome was swabbed from the inside (ID 10).

**Figure 10:**
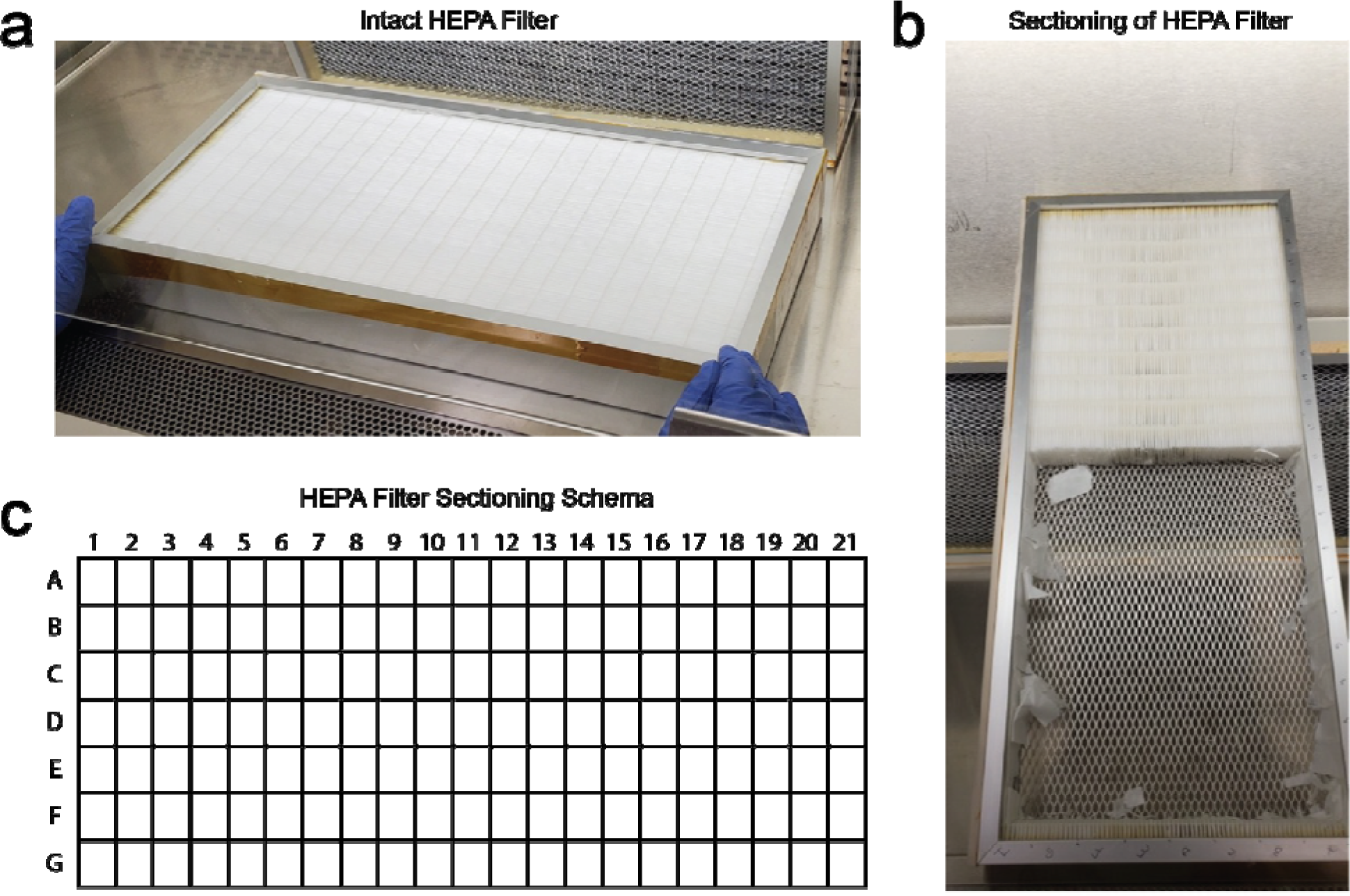
Dragon Capsule HEPA Filter. **(a)** View of the un-cut HEPA filter. **(b)** HEPA filter during sectioning. **(c)** Cutting schema for the HEPA filter. The filter was split into 21 columns and 7 rows, creating a total of 147 preserved sections.

Previous microbial profiling of spacecraft environments has revealed that equipment sterilized on the ground becomes coated in microbial life in space due to interactions with crew and the introduction of equipment that has not undergone sterilization^65^. Subsequent microbial monitoring assays performed on the ISS have detected novel, spaceflight-specific species on the ISS^66^. Once in space, surface microbes are subject to the unique microgravity and radiation environment of flight, which will influence evolutionary trajectory. The potential impact of this influence on pathogenesis is a concern for long-duration space missions, especially given that changes in host-pathogen interactions may also be affected during spaceflight^67^.

## Discussion

We report here on biospecimen samples collected from the SpaceX I4 Mission, the most comprehensive human biological specimen collection effort performed on an astronaut cohort to date. The extensive archive of biospecimens included venous blood, dried blood spot cards, saliva, urine, stool, microbiome body swabs, skin biopsies, and environmental capsule swabs. The study objective was to establish a foundational set of methods for biospecimen collection and banking on commercial spaceflight missions suitable for multi-omic and molecular analysis. Biospecimens were collected to enable comprehensive, multi-omic profiles, which can then be used to develop molecular catalogs with higher resolution of human responses to spaceflight. Select, targeted measures in clinical labs (CLIA) were also performed immediately after sample collection (CBC, CMP), and samples and viable cells were preserved in a long-term Cornell Aerospace Medicine Biobank, such that additional assays and measures can be conducted in the future.

There are several reasons why rigorous biospecimen collection methods for commercial and private spaceflight missions must be developed, which are scalable and translational across populations, missions, and mission parameters. First, little is known about the biological and clinical responses that occur in civilians during and after space travel. While professional astronauts are generally young, healthy, and extensively trained, civilian astronauts have been, and likely will be, far more heterogeneous. They will possess a variety of phenotypes, including older ages, different health backgrounds, and greater medication use, and may experience different medical conditions, risks, and comorbidities. Careful molecular characterization will be beneficial for the development of appropriate baseline metrics and countermeasures and, therefore, beneficial for the individual spaceflight experience. In the future, such analyses may enable precision medicine applications aimed at optimizing countermeasures for each individual astronaut who enters and returns safely from space^68, 69^.

Second, multi-omic studies inherently present a large number of measurements within a small set of subjects. These high-dimensional datasets present numerous potential challenges with regard to amplification of noise, risk of overfitting, and false discoveries^70^. At all times, scientists engaged in multi-omic analyses must take special care that true biological variance is what has been measured. The introduction of experimental variance through the progression from sample collection, transport, storage, to sequencing and analysis can introduce artifacts of variance that render the detection of true biological variance and interpretation of results more difficult. For this reason, tight adherence to experimental controls or annotation at every step of the experimental condition is crucial. Careful annotation allows for the assignment of class variables in post hoc analysis. Among such applications are the attempt to detect batch effects or determine the impact of variations in temperature (collection, storage, or transport)^71^.

The necessary means to address experimental variance are longitudinal sampling and specimen aliquoting. Longitudinal sampling (i.e. collecting numerous serial samples from each test condition) from pre-flight, in-flight, and post-flight allows for greater statistical power when assessing changes attributable to spaceflight. In addition, each sample collected should be divided upon collection into multiple aliquots. This better assures that freeze-thaw cycles can be avoided in the analysis stage, as freeze-thaw events can introduce considerable experimental variance depending on the molecular class being measured. Maintaining all samples at their optimal storage temperature at all times, typically −80°C or lower, is crucial^72^. Special attention must be given to how the collection and storage methods in-flight vary in relation to the conditions on Earth. Spaceflight presents considerable differences in the operating environment, where ground conditions are far easier to control than flight. In practice, this may limit the types of samples that can be collected during flight.

Third, rigorous methods must be developed and followed to pursue comparisons across missions with varying design parameters. In this consideration, there is an argument for the development of specimen collection, transport, storage, processing, analysis, and reporting standards. At the same time, this must be balanced with the flexibility required for innovation since standards can sometimes limit advancement in methodology. In the present study, common methods were used for the Inspiration4 and the forthcoming Polaris Dawn and Axiom missions. However, selected methods may require optimization for Polaris Dawn to increase the yields during sample processing and adapt to unique parameters imposed by the anticipated spacewalk (extravehicular activity; EVA). Moreover, within standards or best practices, unique research for each mission may require alteration of previously successful methods. With these considerations in mind, we must balance methodology standardization with advances in methodology options and mission-specific objectives.

As the commercial spaceflight sector gains momentum and more astronauts with different health profiles and backgrounds have access to space, comprehensive data on the biological impact of short-duration spaceflight is of paramount importance. Such data will further expand our understanding and knowledge of how spaceflight affects human physiology, microbial adaptations, and environmental biology. The use of integrative omics technologies for civilian astronauts will unveil novel data on genomics, proteomics, metabolomics, and transcriptomics. Creating multi-omic datasets from spaceflight studies on astronaut cohorts will further advance our understanding, inform future mission planning, and help discover what appropriate countermeasures can be developed to minimize future risk and enhance performance.

Validating sample collection methodologies initially in short-duration commercial spaceflight is a key step for future human health research in long-duration and exploration-class missions to the Moon and beyond. To help meet these challenges, we have established the SOMA protocols, which detail standard multi-omic measures of astronaut health and protocols for sample collection from astronaut cohorts. Although the all-civilian Inspiration4 crew pioneered the first use of the SOMA protocols, the methodology outlined here is robust and generalizable, making it applicable to future astronaut crews from any commercial mission provider (e.g., SpaceX, Axiom Space, Sierra Space, Blue Origin) or space agencies (NASA, ESA, JAXA, ROSCOSMOS). Furthermore, the SOMA banking, sequencing, and processing methods are a springboard for continuing biospecimen analysis and expanding our knowledge of multi-omic dynamics before, during, and after human spaceflight missions, providing a molecular roadmap for crew health, medical biometrics, and possible countermeasures.

## Acknowledgements

We thank the Scientific Computing Unit (SCU) at WCM and the Genomics, Epigenomics, and Biorepository Cores, the NIH (R01MH117406) and NASA (NNX14AH50G, NNX17AB26G, 80NSSC22K0254, NNH18ZTT001N-FG2, NNX16AO69A, 80NSSC23K0832), the LLS (MCL7001-18, LLS 9238-16, 7029-23), as well as Igor Tulchinsky and the WorldQuant Foundation, the GI Research Foundation (GIRF), the Radvinsky/Chudnovsky family. We thank JJ Hastings for early protocol work. JK thanks MOGAM Science Foundation. We also thank Jennifer Conrad for CPT photography.

## Methods

### Venous Blood Draw

Venipuncture was performed on each subject using a BD Vacutainer® Safety-Lok™ blood collection set (BD Biosciences, #367281) and a Vacutainer one-use holder (BD Biosciences, 364815). The puncture site was located near the cubital fossa and was sterilized with a BZK antiseptic towelette (Dynarex, Reorder No. 1303). Blood was collected into 1 serum separator tube (SST, BD Biosciences: #367987, Lot: #1158449, #1034773), 2 cell processing tubes (CPT, BD Biosciences: #362753, Lot: #1133477, #1012161), 1 blood RNA tube (bRNA, PAXgene: #762165, Lot: #1021333), 1 cell-free DNA BCT (cfDNA BCT, Streck: #230470, Lot: #11530331), and 4 K2 EDTA blood collection tubes (BD Biosciences, #367844, Lot: #0345756) per crew member per time point. For samples collected in Hawthorne, blood was drawn at SpaceX headquarters, then immediately transported to USC for processing. Samples collected at Cape Canaveral were processed on-site.

### Blood Tube Processing

For processing, serum separator tubes (SST) were centrifuged at 1300xg for 10 minutes. 500uL aliquots of serum were aliquoted into 1mL Matrix 2D Screw Tubes (ThermoFisher, 3741-WP1D-BR) and stored at −80°C. SST tubes were recapped and stored at −20°C to preserve the red blood cell pellet.

Cell processing tubes were centrifuged at 1800xg for 30 minutes. Plasma was aliquoted into 1mL Matrix 2D Screw Tubes and stored at −80°C. 5mL of 2% FBS (ThermoFisher, #26140079) in PBS (ThermoFisher, #10010023) was added to the CPT tube to resuspend PBMCs. PBMC suspension was transferred to a clean 15mL conical tube. The total volume was brought to 15mL with 2% FBS in PBS. The tube was centrifuged for 15 minutes at 300xg. Supernatant was discarded. PBMCs were resuspended 6mL of 10% DMSO (Millipore Sigma, #D4540-500mL) in FBS. 1mL of PBMCs were moved to 6 cryogenic vials (Corning, #8672). Cryovials were placed in a Mr. Frosty^TM^ (ThermoFisher, #5100-0001) and stored at −80°C. CPTs were recapped and stored at −20°C to preserve the red blood cell pellet.

Cell-free DNA blood collection tubes (cfDNA BCTs) were centrifuged at 300xg for 20 minutes. Plasma was transferred to a 15mL conical tube. Plasma was centrifuged 5000xg for 10 minutes. 500uL aliquots of plasma were aliquoted into 1mL Matrix 2D Screw Tubes and stored at −80°C. cfDNA BCTs were recapped and stored at −20°C to preserve the red blood cell pellet.

PAXgene blood RNA tubes were processed according to the manufacturer’s instructions. Briefly, tubes were left upright for a minimum of 2 hours before freezing at −20°C. For RNA extraction, tubes were thawed and processed with the PAXgene blood RNA kit (Qiagen, #762164).

### Extracellular Vesicles and Particles (EVPs) Isolation

One 4mL K2 EDTA tube was shipped on ice overnight to WCM for processing. Blood was centrifuged at 500 × g for 10 minutes, then plasma was transferred to a new tube and centrifuged at 3000 × g for 20 minutes, and the supernatant was collected and stored at −80°C for EVP isolation. Plasma volumes ranged between 0.6 - 1.7 ml. Plasma was later thawed for downstream processing, when concentrations were measured. Plasma samples were thawed on ice and EVPs were isolated by sequential ultracentrifugation, as previously described (Hoshino et al., 2020). Briefly, samples were centrifuged at 12,000 × g for 20 minutes to remove microvesicles, then EVPs were collected by ultracentrifugation in a Beckman Coulter Optima XE or XPE ultracentrifuge at 100,000 × g for 70 minutes. EVPs were then washed in PBS and pelleted again by ultracentrifugation at 100,000 × g for 70 minutes. The final EVP pellet was resuspended in PBS.

### Dried Blood Spot (DBS)

Crew members warmed their hands and massaged their finger towards the fingertip to enrich blood flow towards the puncture site. The puncture site was sterilized using a BZK antiseptic towelette (Dynarex, Reorder No. 1303). Skin was punctured using a contact-activated lancet (BD Biosciences, #366593) or a 21-gauge needle (BD Biosciences, #305167), depending on crew member preference. Capillary blood was collected onto the Whatman 903 Protein Saver DBS cards (Cytiva, #10534612). Blood was transferred by touching only the blood droplet to the surface of the DBS card. DBS cards were stored at room temperature with a desiccant pack (Cytiva, #10548239).

### Saliva

To collect crude saliva, crew members uncapped and spit into a sterile, PCR-clean, 5mL screw-cap tube (Eppendorf, 30122330). Crew spit repeatedly until at least 1mL was collected. Saliva was transported to a sterile flow hood and separated into 500uL aliquots. Aliquots were frozen at −80°C. To collect preserved saliva, crew members used the OMNIgene ORAL kit (DNA Genotek, OME-505). Crew members spit into the kit’s tube until they reached the fill line. The tube was re-capped, which released the preservative liquid. Tubes were inverted to mix the saliva and preservative before being placed at −20°C for storage. After all timepoints were collected, DNA, RNA, and protein were extracted using the AllPrep DNA/RNA/Protein kit (Qiagen, #47054). Sample concentrations were measured with Qubit high sensitivity dsDNA and RNA platform. Proteins were quantified with the Pierce™ Rapid Gold BCA Protein Assay Kit (Thermo Scientific, #A53225) on Promega GloMax Plate Reader.

### Urine

Crew members urinated into sterile specimen containers (Thermo Scientific, #13-711-56). The container was stored at 4C until it was prepared for long-term storage. To prepare urine samples for long-term storage, urine was aliquoted into 1mL, 15mL, and 50mL tubes. Half of the urine was immediately placed at −80°C. The other half had urine conditioning buffer (Zymo, #D3061-1-140) added to the sample before placing in the −80°C freezer.

### Stool Collection

Crew members isolated a stool sample using a paper toilet accessory (DNA Genotek, OM-AC1). Stool was transferred into and OMNIgene•GUT tube (DNAgenotek, OMR-200) and an OMNImet•GUT tube (DNA Genotek, ME-200). Tubes were placed at −80°C for long-term storage. For nucleic acid extraction, 200uL of each tube was allocated for DNA extraction with the QIAGEN PowerFecal Pro kit and 200uL was allocated to RNA extraction with the QIAGEN PowerViral kit. The remaining sample was split into 500uL aliquots and re-stored at −80°C.

### Swab Collection

Crew members put on gloves and remove a sterile swab from its packaging. For collection of the postauricular, axillary vault, volar forearm, occiput, umbilicus, gluteal crease, glabella, toe web space, and capsule environment regions, swabs were dipped in nuclease-free water (this step was skipped for oral and nasal swabs) for ground collections. For in-flight collections, HFactor hydrogen infused water was used in place of nuclease-free water. Each body location was swabbed for 30 seconds, using both sides of the swab. Swabs were then placed in 1mL Matrix 2D Screw Tubes containing 400uL of DNA/RNA Shield (Zymo). The tip of the swab was broken off so that only the swab tip was stored in the Matrix 2D Screw Tube. Tubes were stored at 4C.

### Skin Biopsies

Skin biopsies were performed on the deltoid region of the arm. Each site was prepared by application of ChloraPrep and anesthesia was induced with administration of 1% lidocaine with 1:100,000 epinephrine. A trephine punch was used to remove a 3- or 4-mm diameter piece of skin. The resected piece was cut into approximately ⅓ and ⅔ sections. The smaller piece was added to a formalin-filled specimen jar. The larger piece was placed in a cryovial and stored at - 80°C. Surgical defects were closed with 1 or 2 5-0 or 4-0 nylon sutures.

### HEPA Filter

HEPA Filter was taken apart and sectioned under a chemical hood to avoid contamination. The filter contained two parts, an activated carbon component and a HEPA filter. The activated carbon component was discarded and the filter was sectioned using a sterile blade. Sections were placed in individual specimen containers and stored at −20°C.

### Human Subjects Research

All subjects were consented and samples were collected and processed under the approval of the IRB at Weill Cornell Medicine, under Protocol 21-05023569.

### Manuscript Preparation

Figures were generated using Adobe Illustrator and Biorender. Plots were generated in R using ggplot2. SpaceX Dragon capsule images are from the SpaceX Flickr Account (https://www.flickr.com/people/spacex/).

